# The hyperphosphorylation of SARS-CoV-2’s nucleocapsid protein by GSK-3 involves a complex and redundant priming mechanism

**DOI:** 10.1101/2024.05.25.595913

**Authors:** Song Chen, Zunyu He, Jean Kanyo, TuKiet Lam, Grace Ha, Brett Lindenbach, Ya Ha

**Author notes:** These authors contributed equally to this work (Song Chen, Zunyu He).

## Abstract

Upon entry into host cells, the nucleocapsid (N) protein of coronavirus becomes heavily phosphorylated within its central Ser-Arg domain by glycogen synthase kinase-3 (GSK-3). Most substrates for GSK-3 need to be phosphorylated, or primed, by another Ser/Thr kinase four residues downstream of the GSK-3 phosphorylation site. It was widely assumed that the hyperphosphorylation of SARS-CoV-2’s N protein requires independent priming at Ser-188 and Ser-206, which initiates sequential phosphorylation by GSK-3. Here we present biochemical and mass spectrometry data that contradicts this simple model, revealing instead a much more complex and highly cooperative mechanism where redundant priming, as well as exosite docking, enables N protein’s efficient phosphorylation by GSK-3. The R203M and R203K/G204R mutations, found in the recent delta and omicron variants of concern, both hinder N hyperphosphorylation by GSK-3. The mechanistic insights generated in this study suggest a novel approach to treat COVID-19 by combining multiple classes of pharmacological agents to inhibit N hyperphosphorylation.

## Introduction

SARS-CoV-2, the causative agent of the COVID-19 pandemic, is an enveloped positive-sense RNA virus (Hu et al., 2021; Masters, 2006). The virus contains four structural proteins, spike (S), envelope (E), membrane (M) and nucleocapsid (N). One key function of the N protein is to package the viral genome and maintain virion integrity through its multivalent interactions with the single-stranded RNA, oligomerization, and binding to the envelope-embedded M protein (McBride et al., 2014). N comprises of two independently folded domains, the N-terminal domain (NTD) and C-terminal domain (CTD), and three flanking intrinsically disordered regions (IDRs). The amino-terminal half of the central IDR (often referred to as the linker region or LKR) is also called the SR domain due to its many Ser-Arg dipeptide sequences (Fig. 1A). The monomeric NTD, dimeric CTD and positively charged SR domain collectively contribute to RNA binding (Chang et al., 2009; Peng et al., 2020). The CTD dimer also functions as the building block for higher order N oligomers observed inside the virion (Chen et al., 2007; Gui et al., 2017; Klein et al., 2020; Yao et al., 2020). The interaction between N and M likely involves both CTD and the C-terminal IDR (Kuo et al., 2016; Lu et al., 2021).

**Figure 1.**
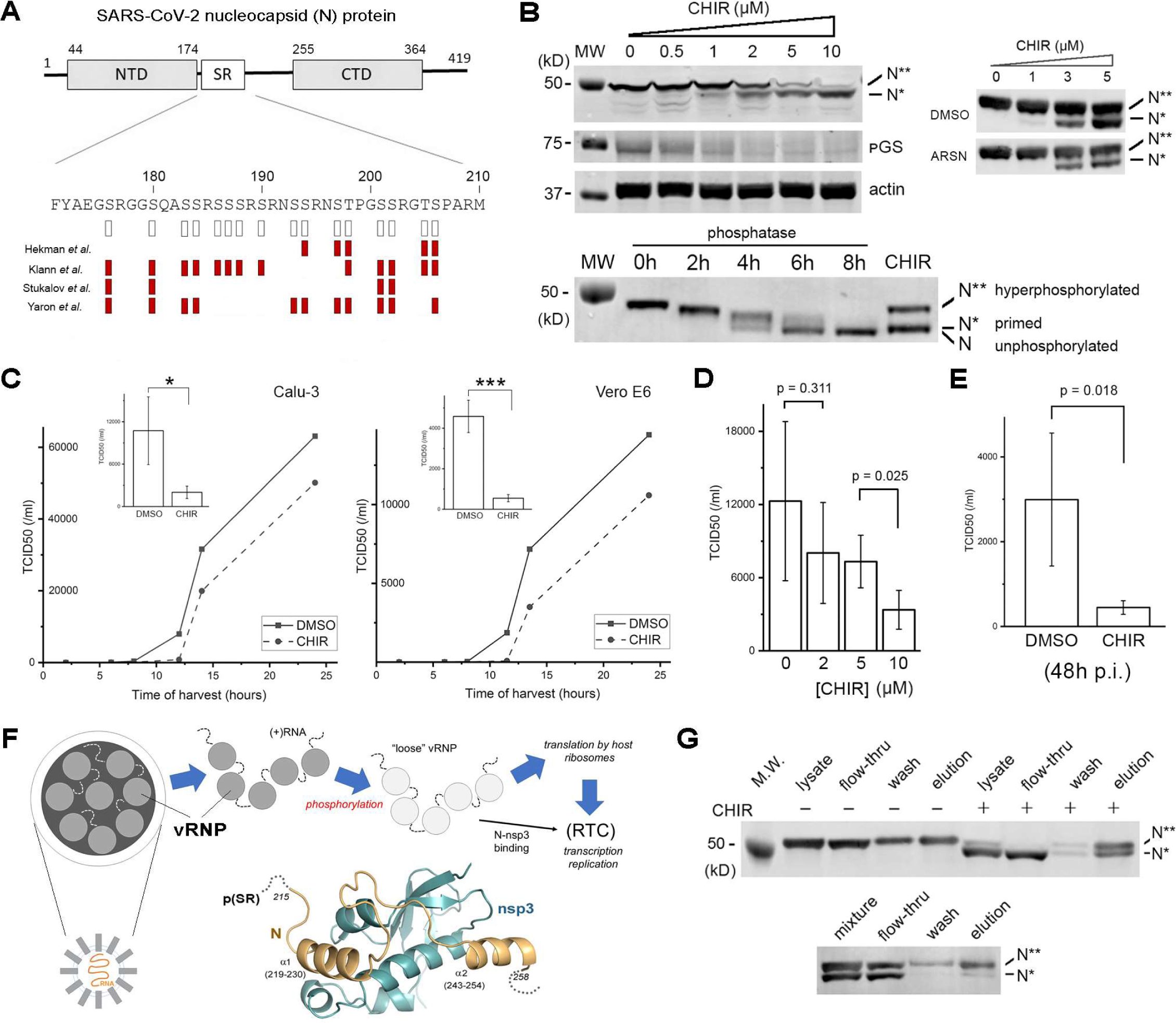
Inhibition of host kinase GSK-3 delays progeny virus release from SARS-CoV-2-infected cells. (**A**) A schematic diagram of the N protein’s domain structure. The 16 potential Ser/Thr phosphorylation sites within the SR domain are indicated by the open bars below the sequence. Phosphorylation sites revealed by phosphoproteomics are indicated by the red bars (only four such studies using infected human cells are shown) (Hekman et al., 2020; Klann et al., 2020; Stukalov et al., 2021; Yaron et al., 2022). (**B**) Supratherapeutic concentration of GSK-3 inhibitor CHIR-99021 (CHIR) is required to suppress N hyperphosphorylation. pGS, glycogen synthase (GS) pSer-641. Untagged N was transiently expressed in 293T cells. (lower panel) No intermediate species between hyperphosphorylated N** and hypophosphorylated N* is observable at 3μM CHIR-99021 (IC_50_ to inhibit N hyperphosphorylation). *In vitro* treatment with calf-intestinal alkaline phosphatase gradually removes phosphate from N and shifts the protein’s electrophoretic mobility from that of the N** to a position slightly lower than that of the N*. (right panel) The 293T cells stably expressing the N construct with a C-terminal (Strep II)_2_-tag were incubated with 500μM arsenite (ARSN) for 30min before CHIR-99021 treatment. (**C**) CHIR-99021 (10μM) alters SARS-CoV-2 replication kinetics in Calu-3 and Vero E6 cells. After cells are briefly inoculated with the virus with a multiplicity of infection (MOI) of 1 and thoroughly washed, culture media are sampled for infectious progeny virus at different time points: the early phase of virus release from infected cells (8-12h) is suppressed by CHIR-99021; the late release phase (>12h) is minimally affected. (Insert) Viral titers at 12h p.i. are compared between DMSO and CHIR-99021 treatment groups (n=3). *, p<0.01; ***, p<0.0001. TCID_50_, medium tissue culture infectious dose. (**D**) Suppression of virus release by CHIR-99021 is dose-dependent. Cultured Calu-3 cells are similarly infected by SARS-CoV-2 and treated with the inhibitor as in (**C**), and viral titers at 12h p.i. are measured and compared (n=3). (**E**) CHIR-99021 (10μM) significantly reduces viral titer 48h after Calu-3 cells are infected by SARS-CoV-2 with a MOI of 0.1 (n=3). (**F**) A schematic diagram illustrating the early steps of SARS-CoV-2’s life cycle. Dashed line denotes the positive-sense and single-stranded viral gRNA. Dark grey circles denote 35∼40 compact viral ribonucleoprotein (vRNP) complexes, or ribonucleosomes, found inside the virion (Klein et al., 2020; Yao et al., 2020). Light grey circles denote less compact ribonucleosomes that can be translated by host ribosomes. In the NMR structure of N (orange) in complex with the ubiquitin-like domain of nsp3 (blue) (PDB accession code: 7PKU) (Bessa et al., 2022), the unresolved SR region is ∼10aa upstream of the α1 helix, and how it influences N:nsp3 binding is not clear. (**G**) The hyperphosphorylated N** has a greater binding affinity for nsp3 than N*. (top) The lysates of 293T cells transiently expressing N (with a C-terminal Strep_2_ tag), with or without CHIR-99021 (10μM) treatment, are passed through a column with immobilized nsp3 ubiquitin-like domain (with a C-terminal Flag_3_ tag) and eluted with free Flag peptide. (bottom) To rule out the possibility that N* could be bound to N** (e.g., through dimerization) in the cell and captured by nsp3 indirectly, cell lysates containing either N** or N* are mixed (1:1 ratio) *in vitro*, passed through the nsp3 column and eluted with Flag peptide.

In addition to its structural role within the virion, the N protein has several crucial functions within the infected cell that are related to its ability to bind and form biomolecular condensates (BMCs) with RNA (Carlson et al., 2020; Cubuk et al., 2021; Iserman et al., 2020; Jack et al., 2021; Lu et al., 2021; Perdikari et al., 2020; Savastano et al., 2020; Wang et al., 2021). After spike-mediated receptor binding and membrane fusion, viral genomic RNA (gRNA), in the form of tightly packed ribonucleoprotein (RNP) complexes, is delivered into the cytosol where it is translated by host ribosomes to yield nonstructural proteins (nsp1-nsp16). A major rearrangement of the N oligomers is required to make the single-stranded RNA sterically accessible to the ribosome (Carlson et al., 2022; Carlson et al., 2020; Dang et al., 2021). After the nsp(s) assemble into the replication transcription complex (RTC) (den Boon et al., 2010), the N-RNA complex is recruited to the membrane-bound supramolecular structure through a direct interaction between N and nsp3 (Cong et al., 2020; Hurst et al., 2013; Hurst et al., 2010; Keane and Giedroc, 2013; Savastano et al., 2020). The transcription of viral RNA involves a unique mechanism called template switching, which enables the production of short subgenomic RNAs (sgRNAs) that guide the synthesis of N and, later, other structural proteins. Some of the newly synthesized N concentrates at the RTC (Scherer et al., 2022), possibly through lipid lipid phase separation (LLPS) (Chau et al., 2023), and regulates the balanced syntheses of short sgRNAs and full-length gRNA (Carlson et al., 2020; Savastano et al., 2020; Verheije Monique et al., 2010; Wu et al., 2014). The final step of the virus’s life cycle involves phase separation of N and gRNA with the membrane-bound M at the nucleocapsid assembly sites and budding of mature virus into the lumen of endoplasmic reticulum-Golgi intermediate compartment (ERGIC) (Lu et al., 2021). Within infected cells, N also interacts with host mRNA (Nabeel-Shah et al., 2022), and a myriad of other factors to modulate host responses (Chen et al., 2021; Gordon et al., 2020a; Gordon et al., 2020b; Min et al., 2023; Zheng et al., 2021; Zhou et al., 2023).

Phosphorylation exerts a large effect on the interaction between N and RNA (Carlson et al., 2022; Carlson et al., 2020; Lu et al., 2021; Maiia et al., 2024; Savastano et al., 2020; Wu et al., 2022). Within the intact virion, a minimally phosphorylated N is probably required to maintain the compact structure of the nucleocapsid (Johnson et al., 2022; Wu et al., 2014; Wu et al., 2009). In virus infected cells, N becomes heavily phosphorylated within the SR domain, which could function as a trigger for conformational change (Carlson et al., 2022; Carlson et al., 2020). Phosphorylation influences template switching as kinase inhibition was found to favor the synthesis of short sgRNAs and suppress the production of full-length gRNA (Wu et al., 2014). The ubiquitous host kinase GSK-3 is responsible for N hyperphosphorylation (Liu et al., 2021; Wu et al., 2009). The presence of multiple GSK-3 phosphorylation series within the central IDR is a conserved feature of all N proteins from the alpha-, beta- and gamma-coronavirus genera (supplementary Fig. 1). Phosphorylation could impact RNA binding directly by neutralizing the highly basic SR domain and thus diminishing its electrostatic interaction with the nucleic acid. The effect could also be indirect as phosphorylation has been shown to strongly influence N’s properties in self-association and phase-separation (Cascarina and Ross, 2022).

Since its initial spillover into the human population in 2019, SARS-CoV-2 has undergone significant adaptive evolution, acquiring mutations especially within the spike protein that enhance receptor binding, membrane fusion or immune evasion (Carabelli et al., 2023; Markov et al., 2023). Intriguingly, besides the notable D614G mutation within the S protein (Volz et al., 2021), the next two most common and positively selected mutations, R203K and G204R, map to the SR domain of the N protein (Harriet et al., 2022; Leary et al., 2021; Rochman et al., 2021; Wu et al., 2021), and are adjacent to Ser-206, one of the two putative priming sites for GSK-3 (Liu et al., 2021; Wu et al., 2009; Yaron et al., 2022). Three variants of concern (VOCs), alpha, gamma, and omicron, all inherited the R203K/G204R double mutation from a parent lineage B.1.1 (Carabelli et al., 2023). The remaining two VOCs also each developed a mutation in this general region (delta: R203M; beta: T205I). In this study we interrogate the biochemical mechanism of N hyperphosphorylation and examine the effect of the three VOC mutations on N phosphorylation. We also explore novel pharmacological strategies to inhibit N hyperphosphorylation with the goal of identifying drug candidates to treat SARS-CoV-2 infection.

## Results and Discussion

### Inhibition of N’s hyperphosphorylation requires supratherapeutic concentrations of GSK-3 inhibitor

When the nucleocapsid (N) protein of SARS-CoV-2 is heterologously expressed in mammalian cells, its SDS-PAGE gel mobility is conspicuously retarded due to heavy phosphorylation (the band labeled with N** in Fig. 1B) (Wu et al., 2009). Host kinase GSK-3 is primarily responsible for this hyperphosphorylation as pharmacological inhibition (Liu et al., 2021; Wu et al., 2009), or genetic ablation (Liu et al., 2021), of the kinase eliminated the N** band. Of the 17 potential GSK-3 sites bearing the (S/T)XXX(S/T) sequence motif (see below), 12 are clustered within the SR domain and solely responsible for the gel shift (Peng et al., 2008; Wu et al., 2009).

As illustrated in Fig. 1B and observed also previously, a higher than usual amount of GSK-3 inhibitor is required to suppress N hyperphosphorylation in cells (Liu et al., 2021; Wu et al., 2009). CHIR-99021, a highly selective and ATP-competitive GSK-3 inhibitor (Bain et al., 2007), has a K_i_ ∼10nM. Given GSK-3’s K_M_ for ATP (26μM), the IC_50_ of CHIR-99021 is calculated to be around 0.8μM based on the Cheng-Prusoff equation *IC_50_ = K_i_ X (1 + [ATP]/K_m,ATP_)* (assuming a cellular ATP concentration of 2mM), which agrees with the observed titration of phosphorylated glycogen synthase (pGS) within the same N-expressing cells. The higher concentration of CHIR-99021, Li^+^ or other GSK-3 inhibitors, required to convert N** to the faster migrating N* suggests that N* is a superb substrate for GSK-3, and its phosphorylation requires less GSK-3 activity than that of glycogen synthase. The observation that no intermediate band is detectable between N** and N* when GSK-3 is partially inhibited also suggests that N hyperphosphorylation is a highly cooperative process (Fig. 1B lower panel).

We examined the effect of phase-separation with RNA on the sensitivity of N hyperphosphorylation to GSK-3 inhibition (Fig. 1B right panel). When expressed in mammalian cells, N has a diffused cytoplasmic distribution but becomes localized to arsenite-induced stress granules, which are mRNA-enriched BMCs (Wang et al., 2021). Arsenite treatment did not lower the IC_50_ of CHIR-99021, suggesting that LLPS itself does not hinder GSK-3-mediated N hyperphosphorylation.

### GSK-3 inhibition delays progeny virus release

Earlier studies using MHV, a model animal coronavirus, implicated GSK-3-mediated N phosphorylation in the readthrough of body transcription regulatory sequence (TRS), which is essential for the syntheses of longer subgenomic mRNAs and the full-length gRNA (Wu et al., 2014; Wu et al., 2009). Surprisingly, under experimental conditions optimized for drug screening (Severson et al., 2007), we found that CHIR-99021 did not protect Vero E6 cells from SARS-CoV-2-induced cytopathic effects (CPE) (data not shown). To better understand the impact of GSK-3 inhibition on viral replication, we infected Calu-3, a human lung epithelial cell line, with SARS-CoV-2 and examined the release of progeny virus into the culture medium. Without CHIR-99021 treatment, new virus started to appear in the medium at 8h post infection (p.i.), and this early phase of viral release is followed by an exponential growth phase starting around ∼12h p.i. (Fig. 1C). GSK-3 inhibition greatly suppressed the early phase of virus release but had little effect on the exponential growth phase. This response is not specific to Calu-3 cells since similar experiment in Vero E6 cells yielded identical results. Using viral tier at 12h p.i. to quantify the compound’s effect, we found that a significant reduction is only achievable at 10μM CHIR-99021 (Fig. 1D). Although these experiments by themselves are not sufficient to prove that the compound’s suppressive effect is mediated through the N protein, the dose-response relationship is consistent with the elevated IC_50_ of CHIR-99021 in blocking N hyperphosphorylation and raises the possibility that complete inhibition of N hyperphosphorylation might be required to delay virus release (Fig. 1B, D). In agreement with a previous report (Liu et al., 2021), we found that 10μM CHIR-99021 suppressed viral titer in the culture medium by >10-fold in an assay where multiple rounds of infection were involved (Fig. 1E).

Upon entering the cytoplasm, phosphorylation of the N protein by host kinases destabilizes the tightly packed nucleocapsid (Carlson et al., 2022; Carlson et al., 2020), which facilitates translation of the positive-sense viral RNA (Fig. 1F). The N protein subsequently plays a role in guiding the single viral genome to the RTC by binding to nsp3 (Cong et al., 2020; Hurst et al., 2013; Hurst et al., 2010; Keane and Giedroc, 2013). To examine the effect of GSK-3-mediated N hyperphosphorylation on nsp3 binding, we passed the cell lysate containing N** and N* through an affinity column with immobilized nsp3 (Fig. 1G). Elution by Flag peptide revealed that the hyperphosphorylated N** outcompeted the hypophosphorylated N* in nsp3 binding, providing direct evidence that phosphorylation of the SR domain could regulate N:nsp3 binding (Koetzner et al., 2022). The observation that the hypophosphorylated N* retains some binding activity toward nsp3 may explain why SARS-CoV-2 can still replicate in the presence of 10μM CHIR-99021 (Fig. 1G). How phosphorylation enhances N:nsp3 binding is unclear at this time as a recent NMR study maps the nsp3-binding segment to residues 219-258, which are slightly downstream of the SR domain (Fig. 1F) (Bessa et al., 2022). It is possible that hyperphosphorylation by GSK-3 affects nsp3 binding indirectly by influencing the accessibility or the confirmation of the 219-258 segment.

Coronavirus is unique among (+)RNA viruses in that its gRNA, when artificially introduced into a cell, has high infectivity only in the presence of the N protein. Using engineered MHV gRNA carrying a luciferase reporter, Koetzner and co-workers recently showed that, without N, luciferase expression was delayed but not completely suppressed (Koetzner et al., 2022). The reason behind the altered kinetics was not fully resolved, but its similarity to what we observed with CHIR-99021 treatment raises the possibility that the essential function of the N protein in supporting viral gRNA infectivity could be compromised by blocking GSK-3-mediated phosphorylation (Fig. 1C).

### N is primed at multiple sites before GSK-3-mediated hyperphosphorylation

Most GSK-3 substrates need to be primed at the i+4 position [(S/T)XXXp(S/T)] by another Ser/Thr kinase [some substrates have multiple (S/T)XXX(S/T) sequence motifs that are joined together, e.g., S^641^…S^645^…S^649^…S^653^ in glycogen synthase (GS), where priming phosphorylation of the most C-terminal (S/T) would enable GSK-3 to phosphorylate the entire series in a sequential manner] (Beurel et al., 2015). The observation that most alpha- and beta-coronavirus N proteins harbor more than one and often long GSK-3 series within their SR domains indicates that hyperphosphorylation could be a functionally important adaptation to mammalian host (supplementary Fig. 1) (Woo et al., 2012).

It was generally believed that the N protein of SARS-CoV-2 contains at least two independent GSK-3 series, each initiated from a separate priming site (Ser-206, Ser-188; highlighted in red in Fig. 2A) (Liu et al., 2021; Wu et al., 2009; Yaron et al., 2022). This hypothesis, however, seems incompatible with the observation that, early during the COVID-19 pandemic, many circulating viruses harbor N mutations that are predicted to disrupt the proposed GSK-3 series: e.g., replacement of Ser-202 by a nonphosphorylatable residue would completely negate the effect of priming at Ser-206 (Fig. 2B; supplementary Table 1). We studied the effect of several of these mutations in 293T cells. As shown in Fig. 2B, none of the three Ser-202 mutations (S202I, S202N, S202R) eliminates N hyperphosphorylation, or renders N hyperphosphorylation more sensitive to CHIR-99021. T205I, a mutation occurring at the C-terminus of another GSK-3 series (series 2 in Fig. 2E), and commonly found in the beta VOC, has no effect either. Interestingly, mutation of Ser-194 (S194L), which is in the same series as Ser-202 but appears in the middle, partially inhibits N hyperphosphorylation, generating a faint N* band in the absence of CHIR-99021. S194L also increases the sensitivity of N hyperphosphorylation to the GSK-3 inhibitor, shifting its IC_50_ closer to the theoretical value (∼1μM).

**Figure 2.**
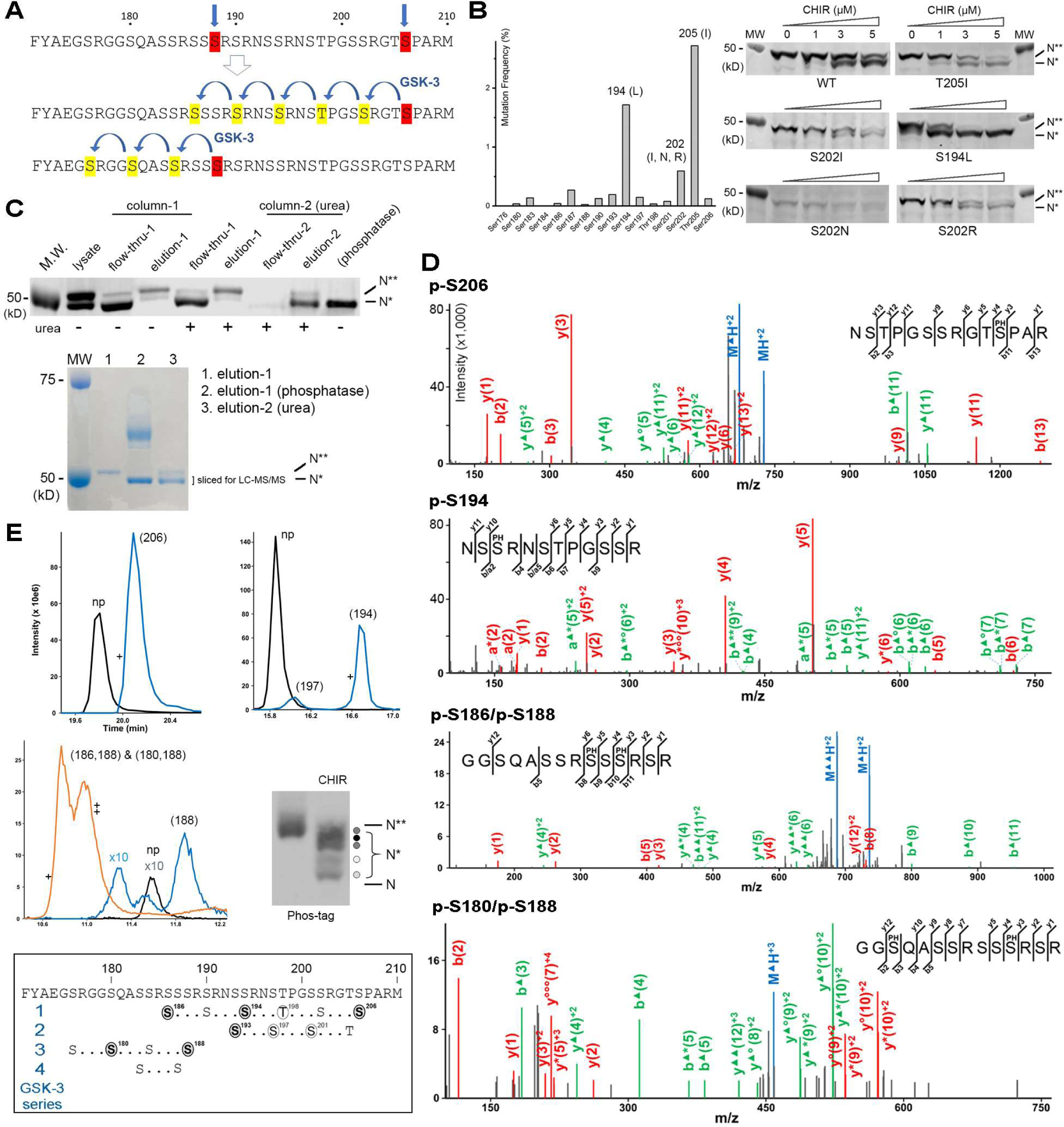
The N protein of SARS-CoV-2 is primed at multiple sites within the SR domain. (**A**) The simple model posits two independent GSK-3 phosphorylation series (highlighted in yellow), each of which requires a separate priming phosphorylation (red). (**B**) Frequency with which the (S/T)s within the SR domain were mutated to a non-phosphorylatable residue during the first two years of the pandemic: the effect of the top five mutations on the sensitivity to CHIR-99021 was tested in 293T cells transiently expression the untagged mutants. All N sequences deposited in the GISAID database at the end of 2021 (∼5,800,000 entries) were downloaded and analyzed (Supplementary Table 1) (Khare et al., 2021). The omicron variant harbors a double mutation within the SR domain (R203K/G204R). After it became dominant after 2021, the frequency of the other mutations within the SR domain was reduced (data not shown). (**C**) Purification of the primed N* for mass spectroscopic analysis. (top) WB with an anti-N antibody; (bottom) Coomassie-stained gel. N with a C-terminal (Strep II)_2_-tag was stably expressed in 293T cells. After treatment with 3μM CHIR-99021, the cell lysate was passed through a StepTactin Sepharose column (column-1), which removed the hyperphosphorylated N**. N* did not bind to the resin, possibly due to aggregation. N* in the flow-through was unfolded by the addition of 6M urea and captured by passing through a second StepTactin column (column-2) in the presence of urea. Urea did not change the gel mobility of either N** or N*. (**D**) MS/MS product ion spectra of the peptide precursor ions used for mapping the phosphorylation sites and quantification. Δ = -98Da (-H_3_PO_4_); O = -18Da (-H_2_O); * = - 17Da (-NH_3_). Monoisotopic ions are colored in blue; product ions with H_3_PO_4_ neutral loss are colored in green; other product ions are colored in red. The precursor ions correspond to singly phosphorylated peptides 196-209 (p-S206) and 192-203 (p-S194) and two different doubly phosphorylated peptides 178-191 (p-S186/p-S188, p-S180/p-S188). (**E**) Estimation of the phosphorylation site occupancy. Elution profiles of the singly phosphorylated (blue) and doubly phosphorylated (orange) peptides, as well as their unphosphorylated counterparts (np; black), from the LC-MS/MS experiment are shown. The + and ++ signs indicate where MS/MS product ion spectra shown in (**D**) were obtained [++ is for the doubly phosphorylated (180, 188) peptide]. Phos-tag gel electrophoresis of 10μM CHIR-99021 treated sample reveals N* is a mixture of several differently phosphorylated species. (boxed) The four GSK-3 phosphorylation series within the SR domain. Majorly phosphorylated sites revealed by LC-MS/MS are indicated by grey circles (occupancy > 30%). Minor sites are indicated by open circles.

To better understand how GSK-3-mediated hyperphosphorylation of the N protein is initiated, we purified the hypophosphorylated N* from a stable 293T cell line expressing a Strep-tagged N (the plasmid was a kind gift from Professor Krogan’s laboratory at UCSF) and characterized it by mass spectroscopy. LC-MS/MS analysis of the purified protein, extracted from the excised Coomassie band and partially digested by trypsin (Fig. 2C), identified a total of 14 phosphorylation sites (supplementary Fig. 2A): three of these are located in the N-terminal IDR (S2, S23, S26), one within the NTD (S105), and ten inside the SR domain (S176, S180, S186, S188, S193, S194, S197, T198, S201, S206) (supplementary Table 2; Fig. 2D; supplementary Fig. 2B; supplementary Fig. 3). Although several previous phosphoproteomic analyses of SARS-CoV-2-infected cells also identified these (and other) sites (Bouhaddou et al., 2020; Davidson et al., 2020; Hekman et al., 2020; Klann et al., 2020; Stukalov et al., 2021; Yaron et al., 2022), the greater amount of the purified protein sample allows us to more reliably determine the occupancy for each phosphorylation site.

The phosphorylation stoichiometry was quantified by comparing the areas under the LC peaks for the phosphorylated peptide (p) and its unphosphorylated counterpart (np), assuming they have similar ionization efficiencies (Fig. 2E; supplementary Table 2). Among the 14 identified sites, only six (S180, S186, S188, S193, S194, S206) are highly occupied, and they are all located within the SR domain (grey circles in Fig. 2E). Ser-188 appears to be fully phosphorylated, and it is primarily found in two doubly phosphorylated peptides: (180, 188) and (186, 188) (the orange peak in Fig. 2E). In contrast, none of the other sites has full occupancy. Ser-206 is found in two singly phosphorylated peptides, but each peptide also has a significant unphosphorylated population (p/np ratio 0.2-2; supplementary Table 2). The possibility that N* could represent an ensemble of several differently phosphorylated species due to partial occupancy at most of the identified sites was tested by running the CHIR-99021-treated sample through a Phos-tag gel, which revealed two groups of bands: the upper group accounts for the majority of N* and may contain up to three species; the lower group contains two minor species (Fig. 2E). Another interesting finding from the mass spectroscopic analysis is that Ser-194, the mutation of which inhibited N hyperphosphorylation (Fig. 2B), is a major site of priming phosphorylation.

The hyperphosphorylated N** was also purified and similarly studied by LC-MS/MS. Unfortunately, none of the identified peptides covers the SR domain, thus leaving open the question about the nature of the hyperphosphorylated state. Our data, however, does reveal that there is no highly occupied phosphorylation site outside the SR domain and thus supports a previous conclusion based on deletion mutagenesis (Peng et al., 2008; Wu et al., 2009).

### The biochemical mechanism of GSK3-mediated hyperphosphorylation

Screening a small collection of kinase inhibitors in our laboratory has yielded two compounds, NCB-0846 and dinaciclib, that weakly suppress N hyperphosphorylation in a GSK-3-independent manner (Fig. 3A). LC-MS/MS analysis of the hypophosphorylated species generated by compound treatment showed that both NCB-0846 and dinaciclib inhibited priming phosphorylation at Ser-188 but had little effect on the phosphorylation of Ser-206 (Fig. 3B). In agreement with their function as priming kinase inhibitors, NCB-0846 and dinaciclib increase the sensitivity of N hyperphosphorylation to GSK-3 inhibitor CHIR-99021 and lithium (Fig. 3C; supplementary Fig. 4).

**Figure 3.**
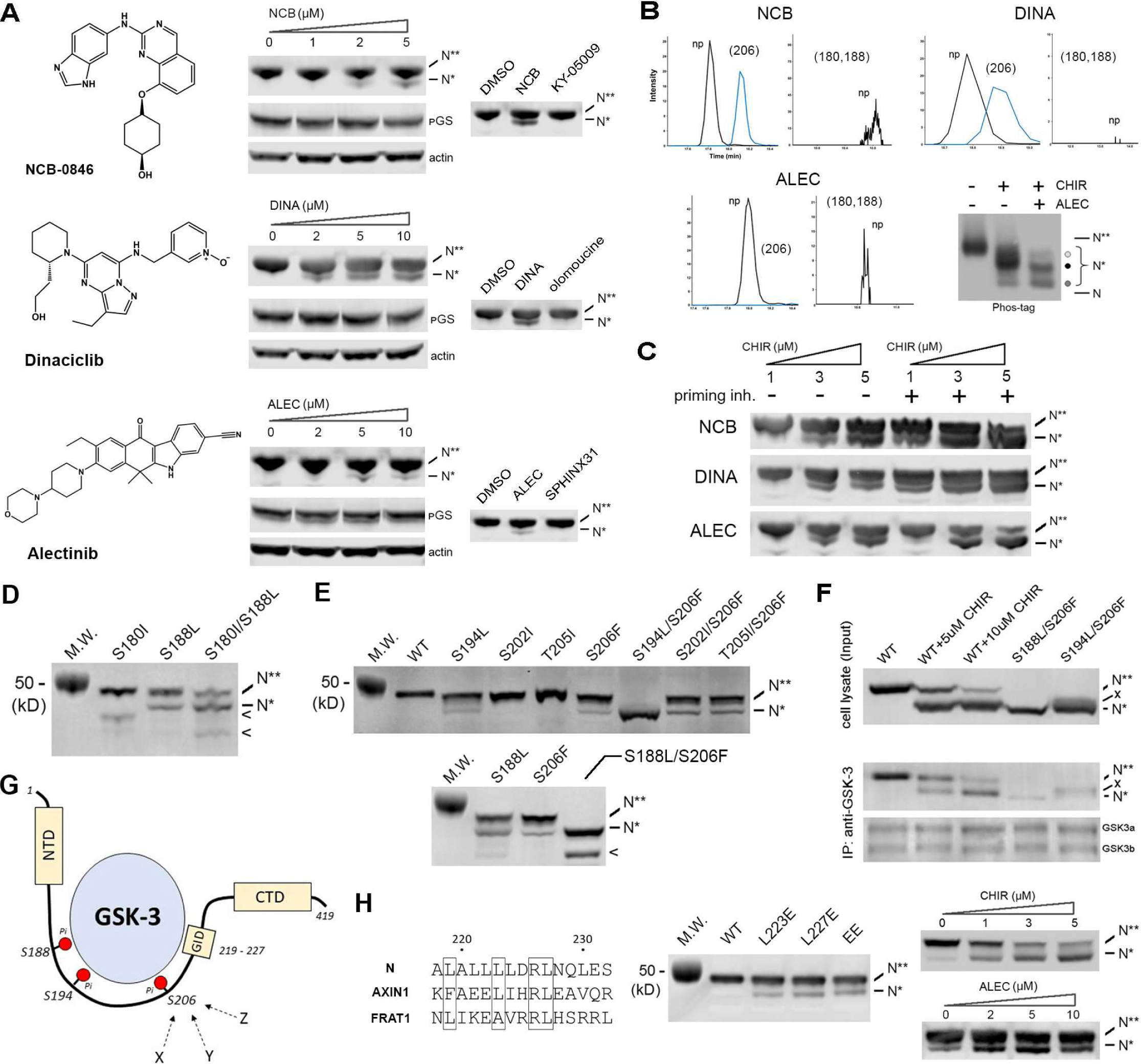
The mechanism of N hyperphosphorylation examined by chemical probes targeting the priming kinase(s) and by mutagenesis. (**A**) NCB-0846, dinaciclib and alectinib weakly inhibit N hyperphosphorylation but do not affect GS phosphorylation (pS641) in 293T cells transiently expressing the N protein with a C-terminal (Strep II)_2_-tag. The effects of NCB-0846 and dinaciclib plateau at 5μM and 10μM, respectively. For reasons yet unclear, the inhibitory activity of alectinib peaks at 5μM and becomes lower at 10μM. At 5μM, TNIK inhibitor KY-05009, CDK2 inhibitor olomoucine and SRPK1 inhibitor SPHINX31 have no effect on N hyperphosphorylation. Inhibitors for CDK1 (Ro-3306), CDK9 (LDC000067), ALK (ceritinib) and CK1δ/ε (PF-670462) also showed no effect (data not shown), suggesting that they are not involved in N hyperphosphorylation. Dinaciclib inhibits CDK1/2/5/9. Since Ser-188 is the only CDK site within the SR domain, and dinaciclib does not reduce its phosphorylation, we can also rule out CDK5 as a priming kinase. (**B**) The faster migrating band generated by NCB-0846, dinaciclib or alectinib treatment was purified and characterized by LC-MS/MS. Elution profiles of the singly phosphorylated (blue) and unphosphorylated peptides (black) containing Ser-206 (NSTPGSSRGTS^206^PAR) and Ser-188 (GGSQASSRSSS^188^RSR) are shown. The two prominent doubly phosphorylated peptides (186, 188) and (180, 188) are missing in all three samples. The effect of combining CHIR-99021 (10μM) and alectinib (10μM) was also examined by Phos-tag gel electrophoresis. (**C**) Priming kinase inhibitor (1μM) increases the sensitivity of N hyperphosphorylation to CHIR-99021 in 293T cells stably expressing the (Strep II)_2_-tagged N. (**D**) The effects of two naturally occurring mutants S188L and S180I from GSK-3 series 3 were examined in 293T cells transiently expressing the untagged N. < indicates a proteolytic fragment. Combining S188L and S180I suppressed N hyperphosphorylation further. (**E**) The effects of three naturally occurring mutations from GSK-3 series 1 (S194L, S202I, S206F) and a mutation (T205I) found in the beta VOC adjacent to Ser-206 were examined. Combining S206F with either S194L or S188L eliminated N hyperphosphorylation. (**F**) The interaction between N and GSK-3 were assessed by co-IP. 293T cells transiently expressing untagged Ns were lysed and immunoprecipitated using an antibody against GSK-3. “x” probably represents a minor species in S194L/S206F where GSK-3 series 3 is phosphorylated. (**G**) A schematic model illustrating multiple contacts between N and GSK-3, which include the interaction of GSK-3 with several phosphorylated S/T(s) and the docking of a GSK-3 interacting domain (GID) slightly downstream of the SR domain (Yun et al., 2022). Priming phosphorylation at each contact site (e.g., Ser-206) is probably catalyzed by more than one kinase. (**H**) Sequence similarity among the GIDs of N, AXIN1 and FRAT1 (Yun et al., 2022). Mutations of the shared (L/F)xx(L/A)xxRL motif partially inhibit but do not eliminate N hyperphosphorylation. The EE double mutation (L223E/L227E) slightly increases the sensitivity to GSK-3 inhibitor CHIR-99021 but has no effect on the sensitivity to priming kinase inhibitor alectinib.

The majority of N remains hyperphosphorylated at concentrations of NCB-0846 or dinaciclib where their effects have plateaued. To address whether this could be due to the chemical probes’ partial inhibition of Ser-188 phosphorylation, we mutated the serine to a non-phosphorylatable amino acid (Fig. 3D). Intriguingly, S188L, a naturally occurring but rare mutation, generates some hypophosphorylated N* instead of converting all the N** to an intermediate species [a result like that produced by S188A (Liu et al., 2021)]. Most of the mutant remains hyperphosphorylated, suggesting that the loss of Ser-188 priming phosphorylation could be overcome by other mechanisms. The absence of any intermediate between N* and N** supports the earlier hypothesis that N hyperphosphorylation is a highly cooperative process. Combining S188L with S180I, which eliminates the (180, 188) double phosphorylation, increases the ratio of N* over N** but does not fundamentally alter the outcome (Fig. 3D). The lower efficacy of the two priming kinase inhibitors is thus likely due to (i) multiple kinases contribute to Ser-188 phosphorylation, and (ii) priming at Ser-188 is not essential for N hyperphosphorylation.

Similar effect on N hyperphosphorylation is observed for another natural mutation, S206F, which eliminated the putative priming site for GSK-3 series 1 (Fig. 2E; Fig. 3E). An unexpected finding, however, is that when S206F is combined with S194L, which eliminated a highly phosphorylated site within the same series, N hyperphosphorylation is abolished. The effect is specific to S194L, as combining S206F with the nearby S202I or T205I does not produce such a drastic reduction (Fig. 3E). Combining S206F with S188L also abolishes hyperphosphorylation (Liu et al., 2021; Wu et al., 2009). Co-IP with GSK-3 reveals that both S206F/S194L and S206F/S188L greatly weaken the binding between N and GSK-3 (Fig. 3F). Taken together, our data suggest a model where GSK-3 engages multiple phosphorylated Ser(s) within the SR domain (Fig. 3G). We hypothesize that these redundant interactions could bypass the classic requirement of GSK-3 for a phosphorylated Ser/Thr at the i+4 position, thus explaining why, for GSK-3 series 1 at least, phosphorylation does not need to occur in a sequential manner (Fig. 2A, B). The Co-IP experiment also shows that the hyperphosphorylated N** binds more tightly to GSK-3, raising the possibility of a positive feedback mechanism where phosphorylation could become more efficient as the reaction proceeds.

Alectinib, the only other known priming kinase inhibitor (Yaron et al., 2022), also suppresses N hyperphosphorylation in 293T cells heterologously expressing the viral protein (Fig. 3A). LC-MS/MS analysis of the faster migrating N* band shows that alectinib inhibited phosphorylation at both Ser-188 and Ser-206 (Fig. 3B). However, unlike S206F/S188L, alectinib only weakly inhibited N hyperphosphorylation, and it is no more efficacious than NCB-0846 or dinaciclib, suggesting again that there exist multiple priming kinases. Phos-tag gel electrophoresis of the sample treated with both CHIR-99021 and alectinib confirms that alectinib is not sufficient to reduce all the hypophosphorylated N* species to a fully dephosphorylated state (Fig. 3B). Future studies are required to identify the kinase targets for NCB-0846, dinaciclib and alectinib responsible for their suppressive effect on N hyperphosphorylation (TNIK, CDK1/2/5/9 and SRPK1, known targets for the chemical probes, are not involved in priming phosphorylation; Fig. 3A).

One more mechanism to strengthen the binding between N and GSK-3 may involve a short segment of N that bears sequence similarity to the GSK-3 interacting domains (GID) of AXIN1 and FRAT1 (Yun et al., 2022). Mutation of the hydrophobic motif (L223E, L227E) partially suppresses N hyperphosphorylation, increases its sensitivity to CHIR-99021, but has no effect on the response to priming kinase inhibitor (Fig. 3H). Nevertheless, the GID appears to play a smaller role in the overall binding (Fig. 3G), since reducing priming phosphorylation alone, e.g., in S206F/S188L, was found to remove most of the co-IP signal (Fig. 3F). It is also intriguing that the GID overlaps completely with the α1 helix that docks onto nsp3 (Fig. 1F) (Bessa et al., 2022), which raises the possibility that nsp3 and GSK-3 may compete in their binding to N.

### R203M and R203K/G204R negatively impact N hyperphosphorylation

R203M and R203K/G204R, when experimentally introduced into the genomic background of wildtype SARS-CoV-2, increase virus fitness and transmissibility (Johnson et al., 2022; Syed et al., 2021; Wu et al., 2021). The biochemical basis for their positive selection, however, remains uncertain (Gussow et al., 2020; Harriet et al., 2022; Johnson et al., 2022; Leary et al., 2021; Muradyan et al., 2024; Reardon et al., 2023; Rochman et al., 2021; Wu et al., 2021). Here we examine how they affect N phosphorylation due to their proximity to Ser-206, a site that we now know is highly phosphorylated by priming kinases and plays an important role in GSK-3 binding and hyperphosphorylation (Fig. 4A-C).

**Figure 4.**
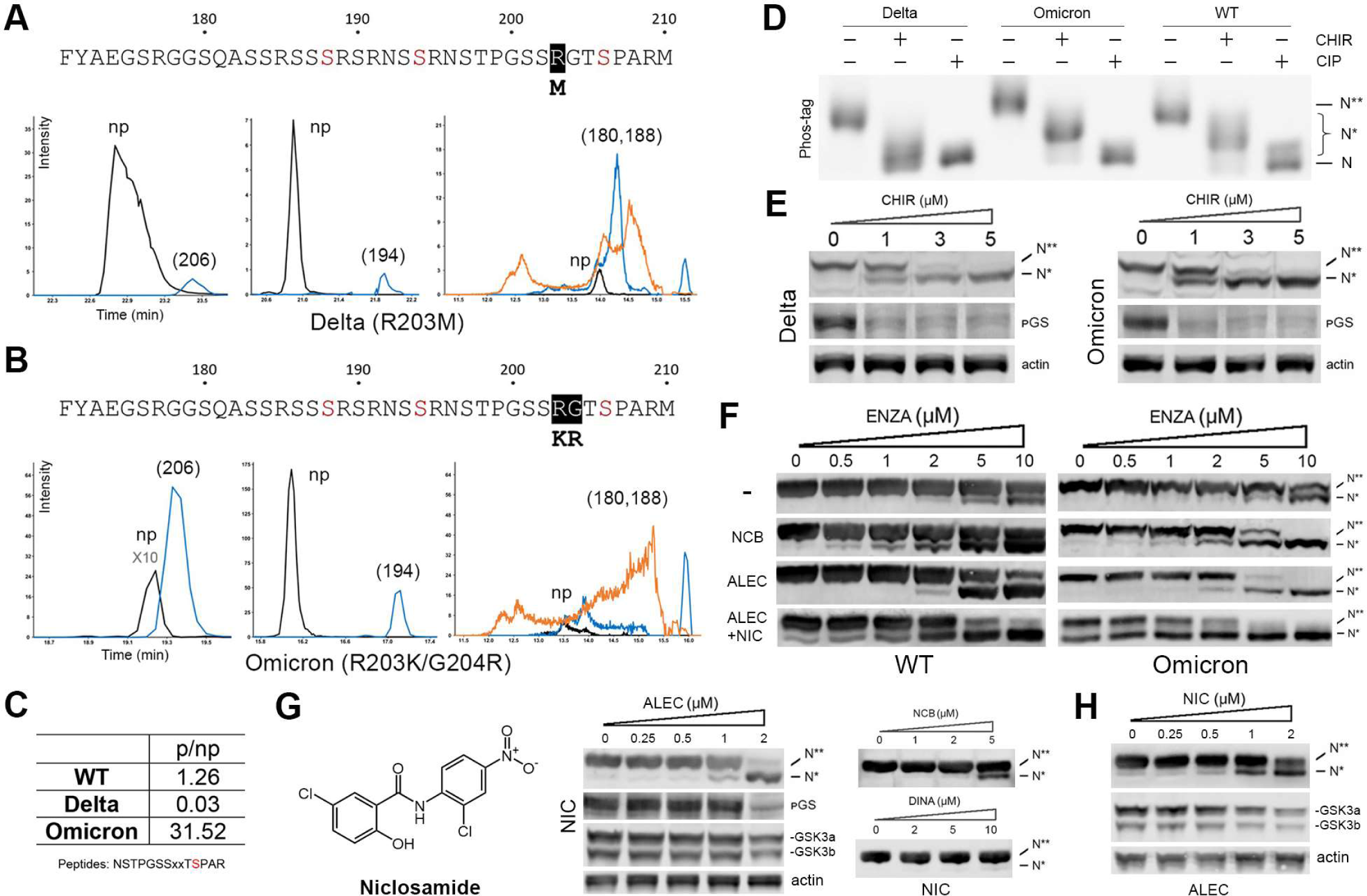
The effect of R203M and R203K/G204R mutations on priming phosphorylation, GSK-3 mediated hyperphosphorylation, and sensitivity to kinase inhibitors. (**A**) R203M, commonly found in the delta VOC, suppresses priming phosphorylation at Ser-206. The hypophosphorylated mutant N*, generated by CHIR-99021 treatment and purified from 293T cells stably expressing the N protein with a C-terminal (Strep II)_2_-tag, was analyzed by LC-MS/MS, and the LC peaks corresponding to phosphopeptides (206), (194) and (180, 188), as well as their unphosphorylated counterparts (np) are shown. (**B**) R203K/G204R of the omicron VOC enhances priming phosphorylation at Ser-206. (**C**) Summary of the mutations’ effect on the ratio between the phosphorylated peptide (p) and its unphosphorylated counterpart (np). “xx” indicates the mutated residues. The WT control was conducted together with the mutants. (**D**) The hypophosphorylated N* species of the delta and omicron variants are analyzed by Phos-tag gel electrophoresis. CIP, calf intestinal alkaline phosphatase. (**E**) Both R203M and R203K/G204R increase the sensitivity of N hyperphosphorylation to CHIR-99021 in 293T cells transiently expressing the untagged mutants. (**F**) Combining 2μM alectinib (ALEC) with 1μM niclosamide (NIC) increases the sensitivity of N hyperphosphorylation to enzastaurin (ENZA) in 293T cells stably expressing the strep-tagged N protein. Enzastaurin is an orally available PKCβ inhibitor but also potently inhibits GSK-3 (Kotliarova et al., 2008; Liu et al., 2021). Although not yet approved by FDA, enzastaurin is well tolerated by patients. The two orally available priming kinase inhibitors (NCB-0846, alectinib) by themselves cannot shift the dose-response curve sufficiently to achieve >50% inhibition of N hyperphosphorylation (WT, omicron) at a physiologically achievable enzastaurin concentration (2μM) (Carducci et al., 2006). (**G**) The synergistic effect between niclosamide (2μM) and alectinib is due to reduced GSK-3 protein level in 293T cells stably expressing the strep-tagged N protein. Niclosamide does not improve the potency or efficacy of the other two priming kinase inhibitors (NCB-0846, dinaciclib). (**H**) In the presence of 2μM alectinib, niclosamide (NIC) suppresses GSK-3 protein level and inhibits N hyperphosphorylation in a dose-dependent manner.

The strep-tagged mutant proteins were expressed in CHIR-99021-treated 293T cells, purified, and characterized by LC-MS/MS (supplementary Fig. 5). Among the six majorly phosphorylated sites, only Ser-206 is significantly affected by the mutations, albeit in opposite ways. R203M greatly reduced Ser-206 phosphorylation, possibly due to the elimination of the upstream (i-3) arginine that is recognized by a basophilic priming kinase (Fig. 4A, C) (Miller and Turk, 2018). Phos-tag gel electrophoresis shows that the primed N*(R203M) has a narrow distribution and migrates closer to the unphosphorylated protein (Fig. 4D). Suppression of Ser-206 priming not only renders N(R203M) hyperphosphorylation more sensitive to GSK-3 inhibitor but also makes the dose-response curve steeper (Fig. 4E; supplementary Fig. 4A, B, D). Furthermore, R203M increases the efficacy of the three priming kinase inhibitors (NCB-0846, dinaciclib, alectinib) (supplementary Fig. 4C). A combinatorial search has identified conditions where N(R203M) hyperphosphorylation can be completely suppressed using orally available kinase inhibitor drugs at concentrations that are physiologically achievable (2μM alectinib plus 2μM enzastaurin; supplementary Fig. 4D) (Carducci et al., 2006; Gadgeel et al., 2014).

In contrast to R203M, R203K/G204R enhanced Ser-206 phosphorylation (Fig. 4B, C). It is possible that introduction of an extra positive charge at the i-2 position makes N a better substrate for the priming kinase, or enables N to be phosphorylated by a new kinase (Miller and Turk, 2018). Intriguingly, increased Ser-206 phosphorylation did not make the protein a better substrate for GSK-3 as previously hypothesized (Johnson et al., 2022). The observation that hyperphosphorylation became more sensitive to GSK-3 inhibitors suggests that the R203K/G204R double mutation itself could negatively impact GSK-3 binding (Fig. 4E, F; supplementary Fig. 4A, B). R203K/G204R did not alter the inhibitory activity of priming kinase inhibitors NCB-0846 and dinaciclib but significantly enhanced the efficacy of alectinib (supplementary Fig. 4C). This is probably because alectinib is the only compound that inhibited Ser-206 phosphorylation, which is expected to exacerbate the negative impact of the double mutation on GSK-3 binding.

It was previously suggested that niclosamide, an anthelmintics drug, could block the binding between N and GSK-3 (Yun et al., 2022). Although our own data do not support this hypothesis, we did observe a small inhibitory effect of niclosamide on N hyperphosphorylation (supplementary Fig. 6A, B). Surprisingly, we discovered that niclosamide greatly enhanced the efficacy and potency of alectinib, while demonstrating little synergistic effect with NCB-0846 or dinaciclib (Fig. 4G). Treating 293T cells with niclosamide and alectinib reduces GSK-3 protein level in a dose-dependent manner, which explains decreased phosphorylation of N and endogenous substrate GS (Fig. 4G, H). It is significant that combining niclosamide (1μM) with alectinib (2μM) improves the potency of GSK-3 inhibitor enzastaurin (Fig. 4F), causing almost complete inhibition at an enzastaurin concentration (2μM) that is therapeutically achievable (Carducci et al., 2006). Although niclosamide has poor bioavailability, sufficient systemic exposure is achievable through oral dosing of a prodrug (Mook et al., 2015). Taken together, our results suggest a novel approach to treat coronavirus infections, especially those caused by the new omicron subvariants harboring the R203K/G204R mutation, using a combination of three orally available drugs to inhibit N hyperphosphorylation. We also noted that the ability of niclosamide/alectinib combination to downregulate GSK-3 protein level could potentially be applied to other disease paradigms, e.g., metabolic, neurological, and neoplastic disorders, where GSK-3 overactivity contributes to pathogenesis (Cohen and Goedert, 2004; McCubrey et al., 2014; Roca and Campillo, 2020).

## Materials and Methods

### Chemical reagents

The following compounds were purchased from Cayman Chemical: CHIR-99021 (CAS#: 252917-06-9), NCB-0846 (CAS#: 1792999-26-8), Dinaciclib (CAS#: 779353-01-4), Alectinib (CAS#: 1256580-46-7), Enzastuarin (CAS#: 170364-57-5), LY2090234 (CAS#: 603288-22-8), KY-05009 (CAS#: 1228280-29-2), Olomoucine (CAS#: 101622-51-9), and SPHINX31 (CAS#: 1818389-84-2). Lithium chloride was obtained from Abcam (ab120853) and sterile filtered for use in cell culture. Niclosamide was obtained from MedChemExpress (CAS#: 50-65-7). Sodium (meta)arsenite was purchased from Chem Cruz (sc-250986).

### Cell culture, transfection, and generation of stable cell line

HEK-293T and VERO E6 cells were purchased from ATCC and cultured in Dulbecco’s Modified Eagle Medium high glucose (Gibco, 11965092) + 10% Fetal Bovine Serum (Gibco, 26140079) + 1% Penicillin/Streptomycin (Gibco, 15140-122). Calu-3 cells were purchased from ATCC and cultured in Eagle’s Minimum Essential Medium (ATCC, 30-2003) with 10% FBS and 1% Penicillin/Streptomycin. Cells were cultured in 37°C incubator with 5% CO2. The tag-free wildtype (Wuhan-hu-1) nucleocapsid protein plasmid was purchased from Addgene (#177937), and the plasmid encoding the Strep-tagged nucleocapsid protein was a kind gift from Dr. Krogan’s lab at UCSF. Transfection of 1 µg per well of plasmids encoding SARS-CoV-2 tag-free nucleocapsid protein was performed using 6 µL per well of Lipofectamine 2000 on HEK-293T cells with 70-80% confluency on 6-well plates (Invitrogen, 11668027). Compounds were added to the cells with fresh media after 6 hours of incubation with plasmids for 24-hour treatment before cell lysis and immunoblotting. HEK-293T cells stably expressing Strep-tagged nucleocapsid protein was generated via lentiviral transduction. 10 μg of the plasmid encoding the Strep-tagged nucleocapsid protein was packaged with 9.183 μg of psPAX2 (Addgene #12260) and 2.768 μg of pVSV-G (Addgene #138479) into lentiviral particles with 50 μL of Lipofectamine 2000 on 80% confluent HEK-293T cells in 6 mL of medium. Lentivirus was harvested once after 48 hours and once after 72 hours by centrifuging the supernatant at 1,000 g for 5 minutes, filtering through 0.45 μm filter (Pall Life Sciences, 12602952) and concentrating to below 500 μL volume with Amicon 100k filter (Millipore, UFC82024). 10,000 HEK293T cells were transduced with 30 μL of concentrated lentivirus in 1.5 mL of medium using 1 µg/ml polybrene (MilliporeSigma™, TR-1003-G). Stable cell line 293T-N was selected and cultured under 1 µg/ml puromycin (Gibco, A1113802). Monoclonal population of the 293T-N was isolated using limiting dilution method. 293T-N cells at 70-80% confluency were treated with compounds for 24 hours to probe N phosphorylation with immunoblotting.

### SARS-CoV-2 viral infection

The infection medium consisted of DMEM (high glucose) supplemented with 2% heat inactivated FBS (Gibco, 16140071), 10 mM HEPES and 1% Penicillin/Streptomycin. The SARS-CoV-2 isolate USA_WA1/2020 (BEI Resources, #NR-52281) was prepared in Dr. Lindenbach’s lab. To generate SARS-CoV-2 viral stock, Vero E6 cells were used for virus propagation. Vero E6 cells were infected with SARS-CoV-2 virus at an MOI of 0.01. After three days, the medium was harvested and centrifuged at 450g for 5 minutes at 4 °C to pellet cell debris. The supernatant was then aliquoted for storage at -80 °C. The SARS-CoV-2 titers in viral stocks were determined in Vero E6 cells by medium tissue culture infectious dose (TCID50) assay. The TCID50 assay were performed on Vero E6 cells using end-point dilution assay. Vero E6 cells were seeded into 96-well plates in growth media and allowed to grow until reaching 80-90% confluence. The growth medium in a 96 well plates was then replaced with 100 μL of 10-fold serial dilutions of the virus in infection medium. After incubation for 5 days in a 5% CO2 incubator, each well was observed under the microscope to determine if cell death has occurred, indicating virus infection. The calculation of TCID50 value was done using the Spearman-Karber method.

To examine the cytopathic effect of SARS-CoV-2, Vero E6 cells were dispensed at 10^4^ cells per well into white 96-well plates (Greiner bio-one, 655074) in infection medium and incubated for 24 hours at 37 °C with 5% CO2. Various concentrations of CHIR-99021 were added to cells and preincubated for 2 hours. Subsequently, the plates were transferred to the BSL3 lab and infected with SARS-CoV-2 virus at a MOI of 0.01. After 3 days of incubation in 5% CO2 incubator, 100 μL of CellTiter-Glo 2.0 Reagent (Promega, G9241) was added to each well. The plates were then shaken for 2 minutes and luminescence was measured using the BioTek Cytation 5 cell imaging multimode reader (Agilent).

For replication kinetics assay, Vero E6 and Calu-3 cells were seeded in 6-well plates 2-3 days prior to infection. Cell counts were conducted on one representative 6-well plate before infection. The cells were then infected with SARS-CoV-2 virus at a MOI of either 1 or 0.1 for 2 hours. Following infection, the cells were washed three times with infection medium and subsequently replenished with infection medium containing CHIR-99021. The cells were maintained in a 5% CO2 incubator and medium samples were collected at different time points. These samples were centrifuged at 450g for 5 minutes and the supernatant was utilized for TCID50 assay.

All work with live SARS-CoV-2 virus was performed in the Biosafety Level 3 laboratory and approved by the Yale University Biosafety Committee.

### SARS-CoV-2 nucleocapsid protein purification

Strep-tagged SARS-CoV-2 nucleocapsid protein with original Wuhan-hu-1 sequence was stably expressed in lentiviral-transduced cell line 293T-N. Cells were treated 24 hours after plating on 100 mm dish by 5 µM of compounds for 24 hours. Buffer A was used to lyse the cells for 15 minutes on ice and cell pellet was removed by centrifugation at max speed for 30 minutes. 40 µL of Strep-Tactin®XT 4Flow high capacity resin (IBA Lifesciences, 2-5030-002) was equilibrated by 200 µL of buffer A for 3 times in an Eppendorf tube and incubated with cell lysate supernatant overnight at 4°C. The resin was spun down at 500 x g for 2 to 5 minutes. 100 µL of buffer B was used to wash the protein-bound resin 5 times before eluting (E1) the hyper-phosphorylated portion of the Nucleocapsid protein with 50 µL of buffer C containing Biotin (IBA Lifesciences, 2-1016-002). The flow-through sample containing the hypo-phosphorylated portion was added 6 M urea to enhance resin binding. 30 µL of fresh Strep Tag II resin was equilibrated by 200 µL of buffer D for 3 times again and incubated with urea-treated flow through sample overnight at 4°C. 100 µL of buffer E was used to wash the protein-bound resin 5 times before eluting the hypo-phosphorylated portion of the Nucleocapsid protein with 40 µL of buffer F once after 24 h and once after 48 hours (E2). Compositions of all buffers used in purification process is listed below:

Buffer A: 25 mM Tris-HCl pH 8.0, 150 mM NaCl, 2 mM EDTA pH 8.0, 0.05% SDS, 1% NP-40

Buffer B: 25 mM Tris-HCl pH 8.0, 500 mM NaCl, 2 mM EDTA pH 8.0, 1% NP-40

Buffer C: 100 mM Tris-HCl pH 8.0, 150 mM NaCl, 1 mM EDTA pH 8.0, 50 mM Biotin

Buffer D: 25 mM Tris-HCl pH 8.0, 150 mM NaCl, 2 mM EDTA pH 8.0, 0.05% SDS, 1% NP-40, 6 M Urea

Buffer E: 25 mM Tris-HCl pH 8.0, 500 mM NaCl, 2 mM EDTA pH 8.0, 1% NP-40, 6 M Urea

Buffer F: 100 mM Tris-HCl pH 8.0, 150 mM NaCl, 1 mM EDTA pH 8.0, 50 mM Biotin, 6 M Urea

Gel electrophoresis was performed on E2 samples with NuPAGE 10% Bis-Tris protein gels (Invitrogen, NP0301) by running 7 µL of sample in each well at 150 V for 4.5 h on ice. Gels were fixed, stained and destained by following standard protocol of Coomassie Brilliant Blue G 250 Staining (SERVA Electrophoresis, 17524). After washing the gels with distilled water, the lower band (hypophosphorylated) of the N* proteins from each lane was cut and submitted for LC-MS/MS analysis.

### LC-MS/MS

Gel slices were cut into small pieces and washed with 1 mL water on a tilt-table for 10 minutes, then washed with 1 mL 50% acetonitrile (ACN)/100 mM NH_4_HCO_3_ (ammonium bicarbonate, ABC) twice for 20 minutes. The samples were reduced by the addition of 100 µL 4.5 mM dithiothreitol (DTT) in 100 mM ABC with incubation at 37°C for 20 minutes. The DTT solution was removed, and the samples were cooled to room temperature. The samples were alkylated by the addition of 100 µL 10 mM iodoacetamide (IAN) in 100 mM ABC with incubation at room temperature in the dark for 20 minutes. The IAN solution was removed, and the gels were washed with 1 ml 50% ACN/100 mM ABC for 20 minutes, then washed with 1 ml 50% ACN/25 mM ABC for 20 minutes. The gels were briefly dried by SpeedVac, then resuspended in 1 gel volume of 25 mM ABC containing 5 ng/µL of digestion grade trypsin (Promega, V5111) and incubated at 37°C for 16 hours for a full digestion or for 15 minutes for a partial digestion, with proteolysis halted by the addition of trifluoroacetic acid (TFA) to a final concentration of 1%. The supernatants containing the tryptic peptides were transferred to new Eppendorf tubes. Residual peptides in the gel bands were extracted with 300 µl 80% ACN/0.1% trifluoroacetic acid (TFA) for 15 minutes, then combined with the original digests and dried in a SpeedVac. Peptides were dissolved in 24 µL MS loading buffer (2% acetonitrile, 0.2% trifluoroacetic acid), with 5 µl injected for LC-MS/MS analysis.

LC-MS/MS analysis was performed on a Thermo Scientific Q Exactive Plus equipped with a Waters nanoAcquity UPLC system utilizing a binary solvent system (A: 100% water, 0.1% formic acid; B: 100% acetonitrile, 0.1% formic acid). Trapping was performed at 5µl/min, 99.5% Buffer A for 3 minutes using a Waters Symmetry® C18 180µm x 20mm trap column. Peptides were separated using an ACQUITY UPLC PST (BEH) C18 nanoACQUITY Column 1.7 µm, 75 µm x 250 mm (37°C) and eluted at 300 nl/min with the following gradient: 3% buffer B at initial conditions; 5% B at 1 minute; 25% B at 45 minutes; 50% B at 65 minutes; 90% B at 70 minutes; 90% B at 75 min; return to initial conditions at 77 minutes. MS was acquired in profile mode over the 300-1,700 m/z range using 1 microscan, 70,000 resolution, AGC target of 3E6, and a maximum injection time of 45 ms. Data dependent MS/MS were acquired in centroid mode on the top 20 precursors per MS scan using 1 microscan, 17,500 resolution, AGC target of 1E5, maximum injection time of 100 ms, and an isolation window of 1.7 m/z. Precursors were fragmented by HCD activation with a collision energy of 28%. MS/MS were collected on species with an intensity threshold of 1E4, charge states 2-6, and peptide match preferred. Dynamic exclusion was set to 20 seconds. A few preliminary runs were collected with a slightly different gradient and +1 species included.

Data was analyzed using Proteome Discoverer (version 2.5) software (Thermo Scientific) and searched in-house using the Mascot algorithm (version 2.8.0) (Matrix Science). The data was searched against the Swissprotein database with taxonomy restricted to Homo sapiens (20,377 sequences) along with a custom database containing the SARS-CoV-2-Nucleocapsid Protein WT and mutant sequences of interest. Search parameters used were trypsin digestion with up to 6 missed cleavages; peptide mass tolerance of 10 ppm; MS/MS fragment tolerance of +0.02 Da; and variable modifications of methionine oxidation, carbamidomethyl cysteine, phosphorylation on serine, threonine, or tyrosine, and deamidation on asparagine and glutamine. Normal and deco database searches were searched, with the confidence level set to 95% (p<0.05). Putative matches to phosphorylated peptides of interest were manually inspected. LC elution peaks of peptides were generated using Xcalibur Qual Browser based on corresponding m/z.

### Immunoblotting

Cells were washed with ice-cold 1x DPBS buffer and then lysed using RIPA buffer (50 mM Tris-HCl pH 7.4, 150 mM NaCl, 1% NP-40, 0.1% SDS, 0.5% DOC, 1 mM EDTA, protease and phosphatase inhibitor cocktail). After centrifugation at max speed for 30 min, samples were collected for BCA assay (Thermo scientific, 23225) to estimate the total protein concentration of each sample. To maximally dissolve hypophosphorylated Nucleocapsid protein in gel loading samples, cell lysates were directly mixed with LDS sample buffer and 1 mM Dithiothreitol (DTT) and was boiled at 90 °C before centrifugation at max speed. Large chunks of precipitates were removed then from the cell lysate samples before running the samples through NuPAGE 10% Bis-Tris protein gels (Invitrogen, NP0301). For blotting phosphatase-treated Nucleocapsid proteins, cells were lysed using a buffer containing 50 mM Tris-HCl pH 7.4, 150 mM NaCl, 1% Triton X-100, 1 mM EDTA, protease inhibitor, and protein samples were treated with 50-100 units of Quick CIP phosphatase (New England Biolabs, M0525S) overnight for complete dephosphorylation. All electrophoresis were run at 150 V for 4 to 5 hours on ice until the 37 kDa protein ladder ran to the end of gel to resolve differently phosphorylated forms of the Nucleocapsid protein. Phospho-tag gels (7.5%, 17well) were purchased from FUJIFILM (192-18001) and run in the same way to resolve differently primed N* proteins. Immunoblotting was then performed with 0.45μm Immobilon-FL PVDF membranes (Millipore, IPFL00010) which were blocked by 5% non-fat milk (americanBIO, AB10109-01000) and then incubated overnight at 4°C with the following primary antibodies: Nucleocapsid protein (Cell Signaling, #33717), Strep-tagged Nucleocapsid protein (Invitrogen, MA5-37747), phospho-Glycogen Synthase (Cell Signaling, 3891S), β-actin (Cell Signaling, 8457S/3700S), GSK-3α/β (Cell Signaling, 5676S). Secondary fluorescent antibodies (LI-COR Biosciences, D10825-15/D10831-15) were used for visualization and imaging by an Odyssey CLx system.

### NSP3 Pull-down

The sequence encoding the NSP3 ubiquitin-like domain 1 (Ubl1, residues 1-110) of SARS-CoV-2 was synthesized by Genscript Biotech and inserted into the pCMV-3Tag-3A plasmid, which contains a C-terminal 3xFlag tag. The construct, Flag-fused NSP3(1-110) plasmid, was transfected into 293T cell using Lipofectamine 2000 and the protein was expressed for 1-2 days. Subsequently, all cells were lysed in lysis buffer (50 mM Tris-HCl, pH 7.4, 150 mM NaCl, 1 mM EDTA, 1% (w/v) Triton X-100 and 5% (w/v) glycerol), supplemented with protease and phosphatase inhibitors.

The NSP3(1-110) protein was then immobilized on Anti-Flag M2 Affinity gel (Sigma Aldrich, A2220), and the cell lysates from stable cell line 293T-N, with or without CHIR-99021 treatment, were passed through the gel. Following this step, the gel was washed with 10 column volumes of lysis buffer and subsequently eluted with 100 µg/mL free Flag peptide (Sigma Aldrich, F3290) in lysis buffer. Finally, all samples underwent immunoblotting analysis.

### GSK3 Co-IP

WT nucleocapsid protein and other nucleocapsid mutant proteins were transiently expressed for 24 hours in 293T cell. All cells were harvested and lysed in lysis buffer (50 mM Tris-HCl, pH 7.4, 150 mM NaCl, 1 mM EDTA, 1% (w/v) Triton X-100 and 5% (w/v) glycerol) supplemented with protease and phosphatase inhibitors. 200 μL of each cell lysate at a total protein concentration of 1 mg/mL was incubated with 4 μL of GSK3α/β antibody overnight at 4`°`C. Following incubation, 80 μL of 25% bead slurry Protein A MagBeads (Genscript Biotech, L00273) was added and incubated with rotation for 3 hours at 4`°`C. The MagBeads were washed five times with 500 μL of cell lysis buffer using a magnetic separation rack. Finally, samples were eluted in 100 μL of 1x SDS sample buffer and boiled for 5 minutes. The eluted samples were then analyzed by SDS-PAGE followed by immunoblotting.

### DNA Cloning

Mutations of phosphorylation sites within the SR region were generated via site-directed mutagenesis based on either the tag-free SARS-CoV-2 Nucleocapsid protein plasmid (Addgene, #177937) or the Strep-tagged SARS-CoV-2 Nucleocapsid protein plasmid (Dr. Krogan’s lab, UCSF). PCR was performed with PrimeSTAR® GXL DNA Polymerase (Takara, R050A), PrimeSTAR GXL Buffer (Takara, SD1967), and dNTP Mix (Takara, SD7178). 50 µL of PCR product was digested by 1μL of DpnI (NEB) for 2 hours. PCR products were purified using DNA gel extraction with NucleoSpin® Gel and PCR Clean-up kit (MACHEREY-NAGEL, 740609.10). Transformation was performed with purified PCR products in Stellar Competent Cells (Takara, 636763). Primers used for PCR and sequencing are listed below:

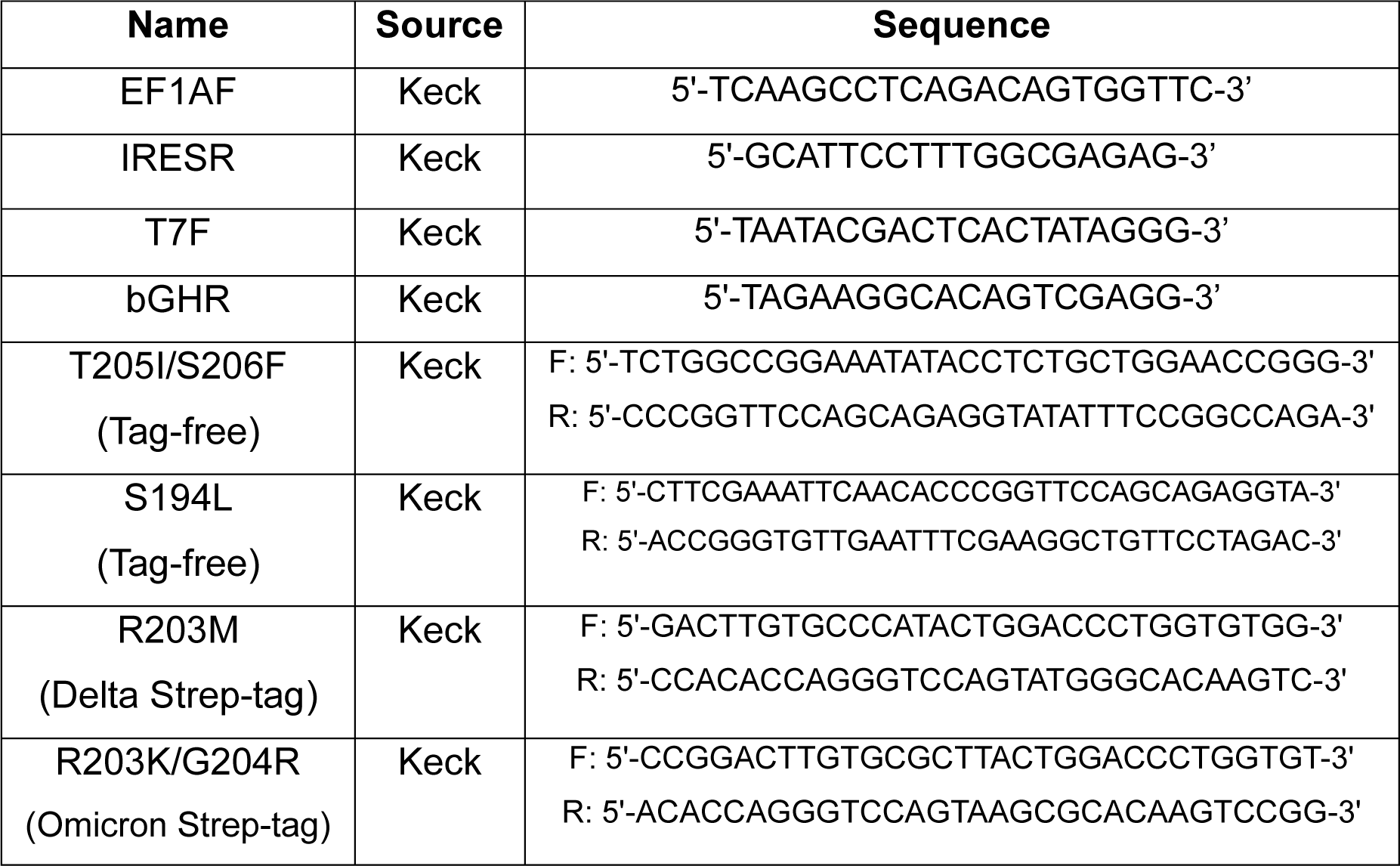
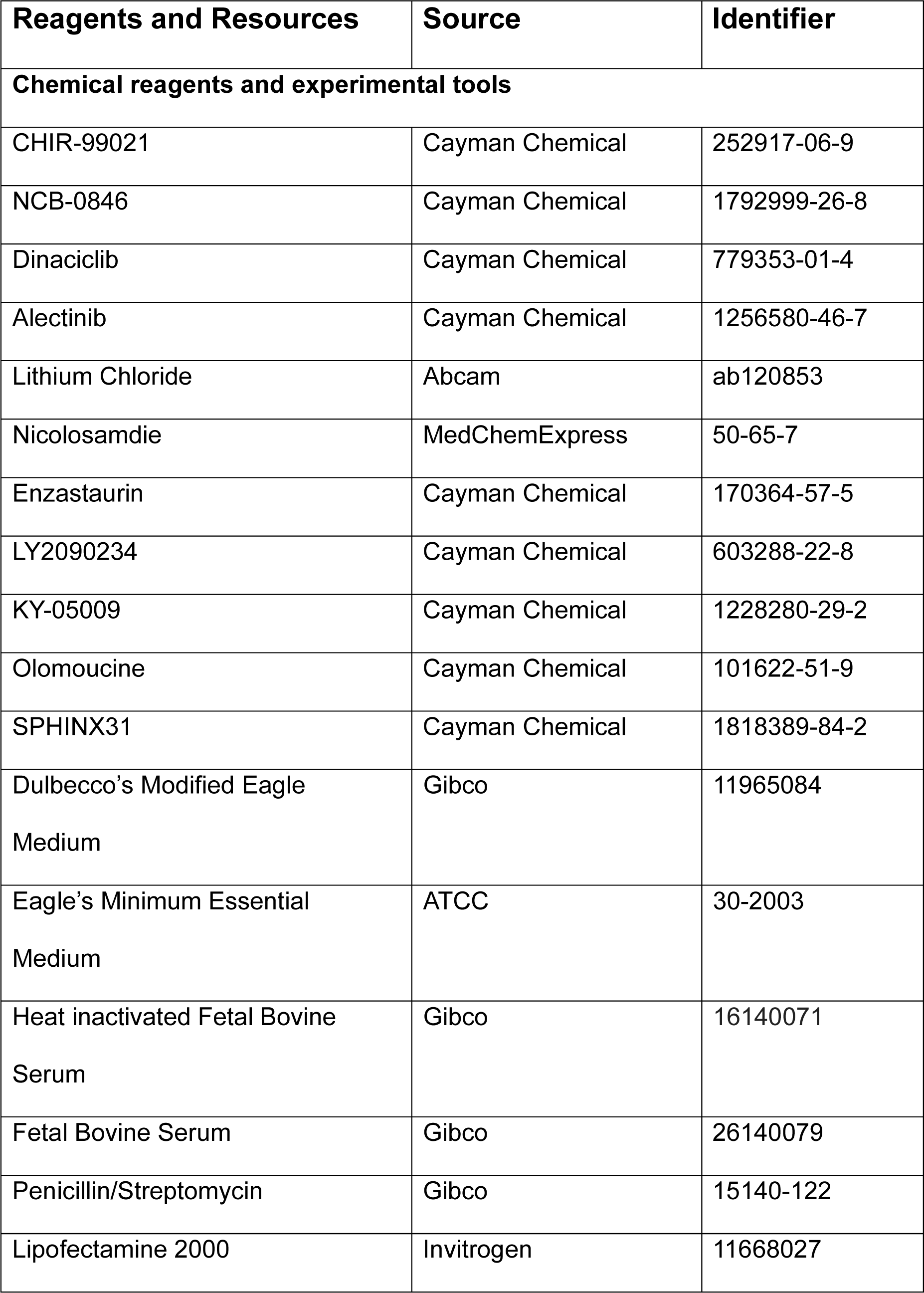
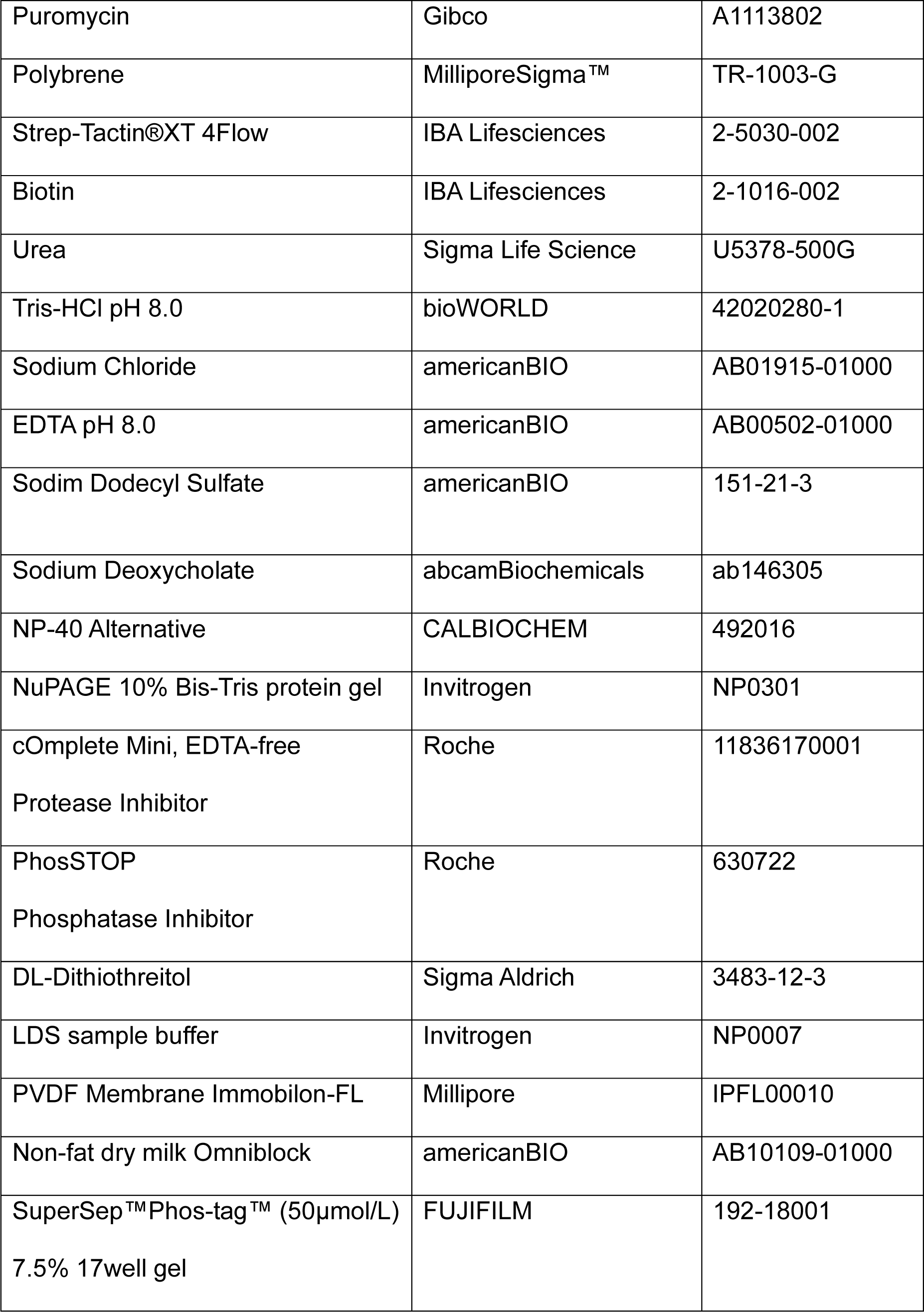
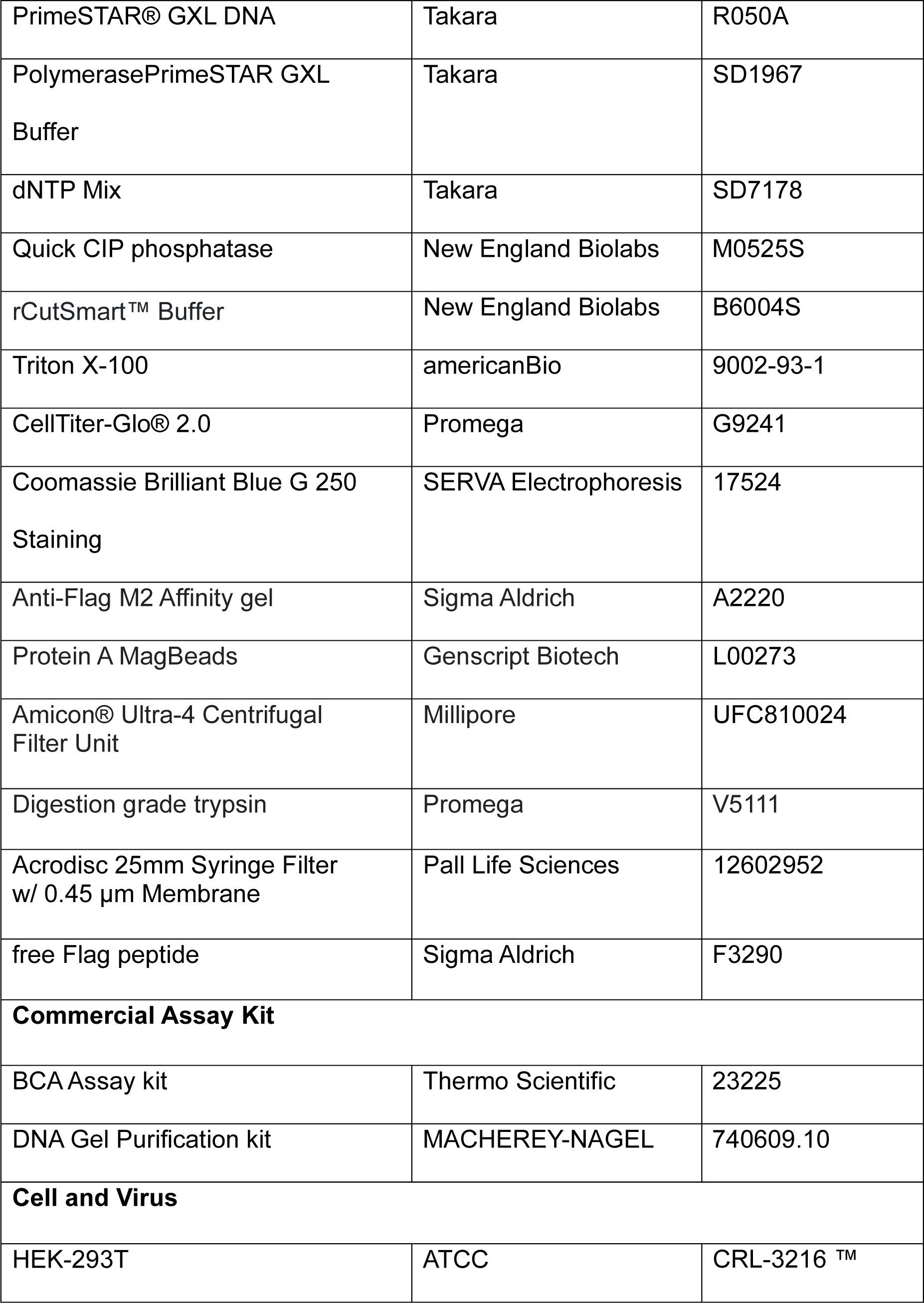
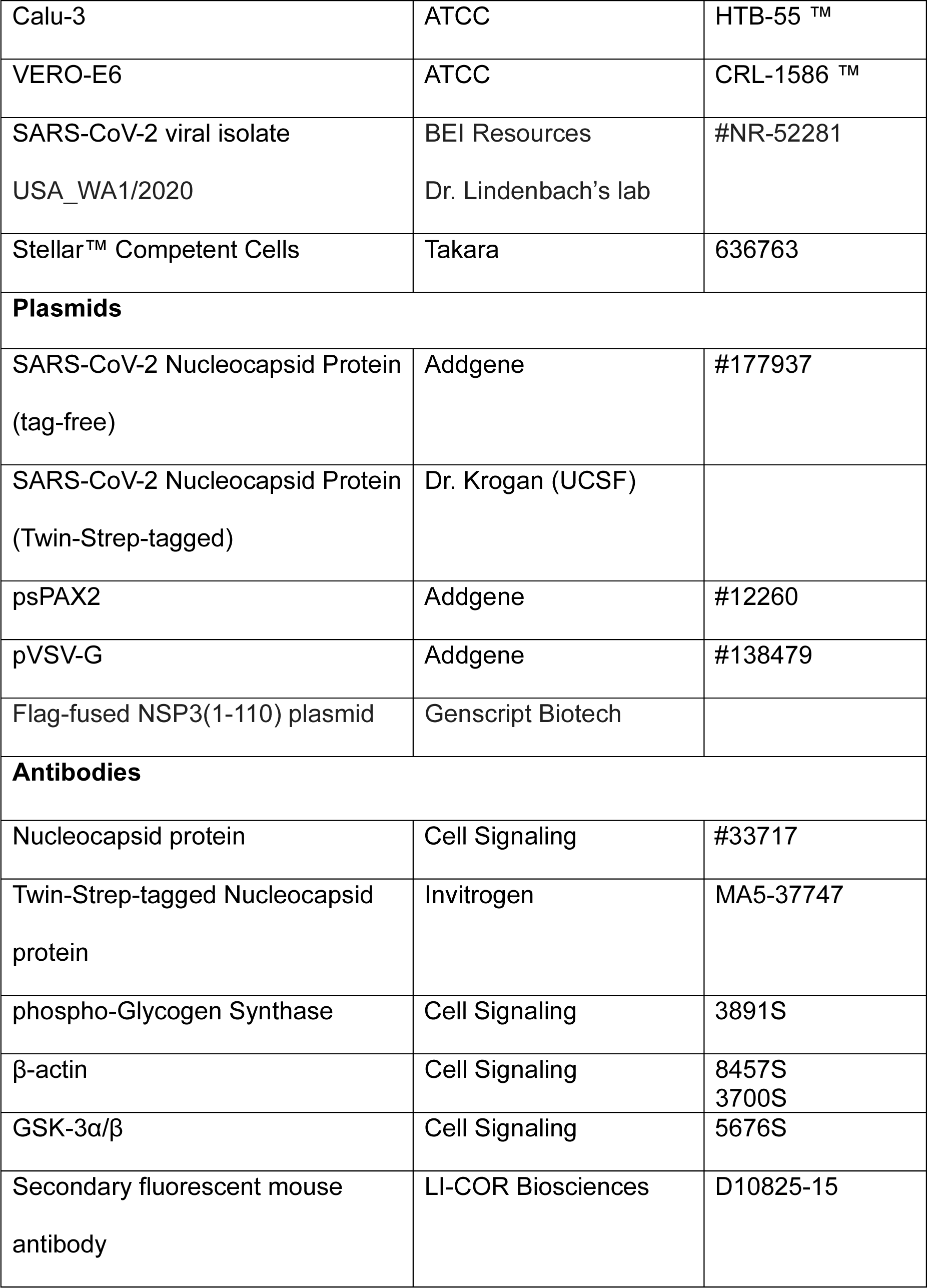
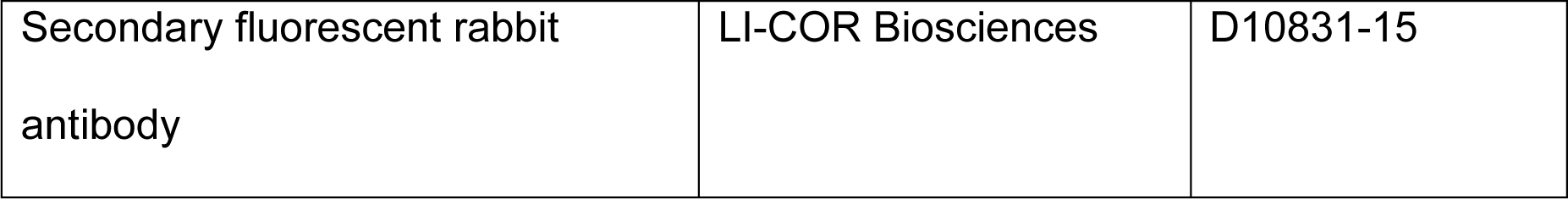

## ACKNOWLEDGEMENTS

We thank Nevan Krogan (UCSF) for sharing the plasmid encoding the Strep-tagged nucleocapsid protein. We thank Florine Collin for her assistance in the mass spectrometry experiment. We thank Benjamin Fontes, Spasov Krasimir, Maren Schniederberend and Anthony D’Abramo for their training and help in the live virus experiment within Yale’s BSL-3 laboratories. We thank Gary Rudnick for sharing his imager and Karen Anderson for sharing her plate reader. This work was partly supported by Yale University (Y.H.), National Institutes of Health grants GM138722 and GM150502 (to Y.H.).

## Supplementary figure legends

**Figure S1.**
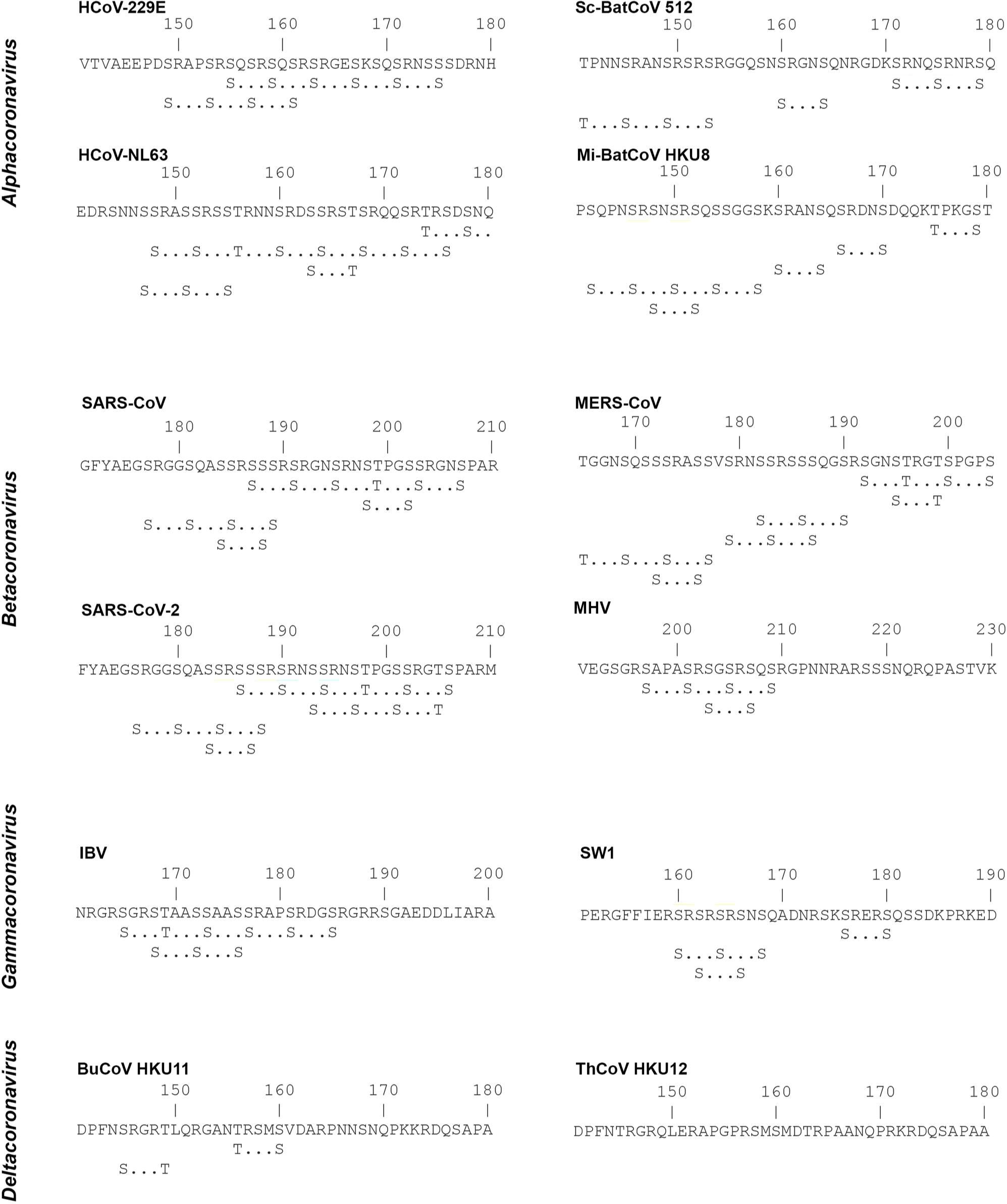
Potential GSK-3 phosphorylation sites within the central IDR. Alpha- and beta-coronaviruses evolved from an ancestral bat virus, whereas gamma- and delta-coronaviruses had an avian origin (Woo et al., 2012). Some deltacoronavirus N proteins do not carry any (S/T)XXX(S/T) sequence motif or have an apparent SR domain.

**Figure S2.**
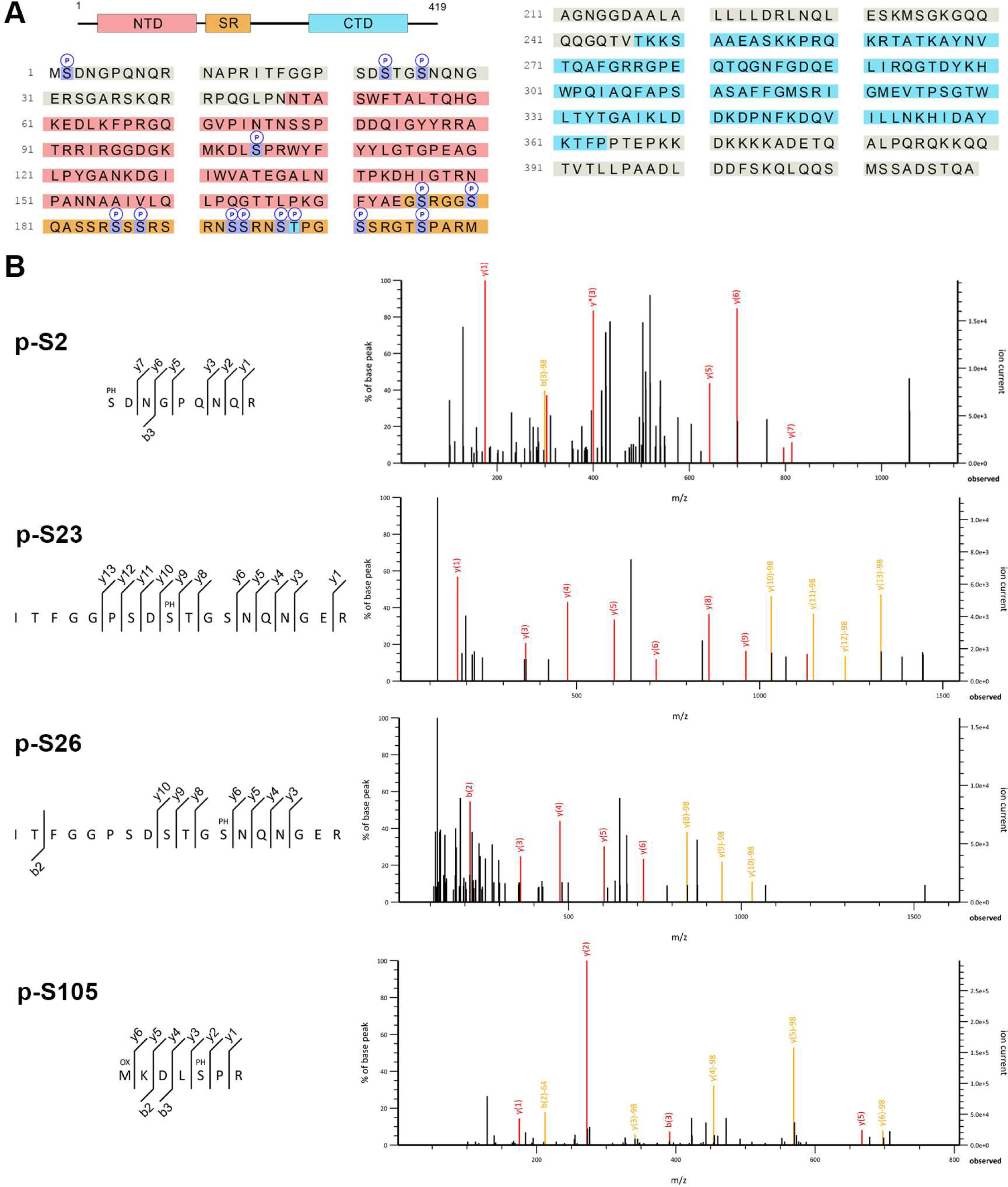
Phosphorylation sites detected by LC-MS/MS. (**A**) The entire sequence of N is covered by peptides generated through partial tryptic digestion and identified by LC-MS/MS (the C-terminal Strep tag is also covered but not shown). Resolved phosphorylation sites are indicated. (**B**) MS/MS product ion spectra of the peptide precursor ions used for mapping the phosphorylation sites outside the SR domain.

**Figure S3.**
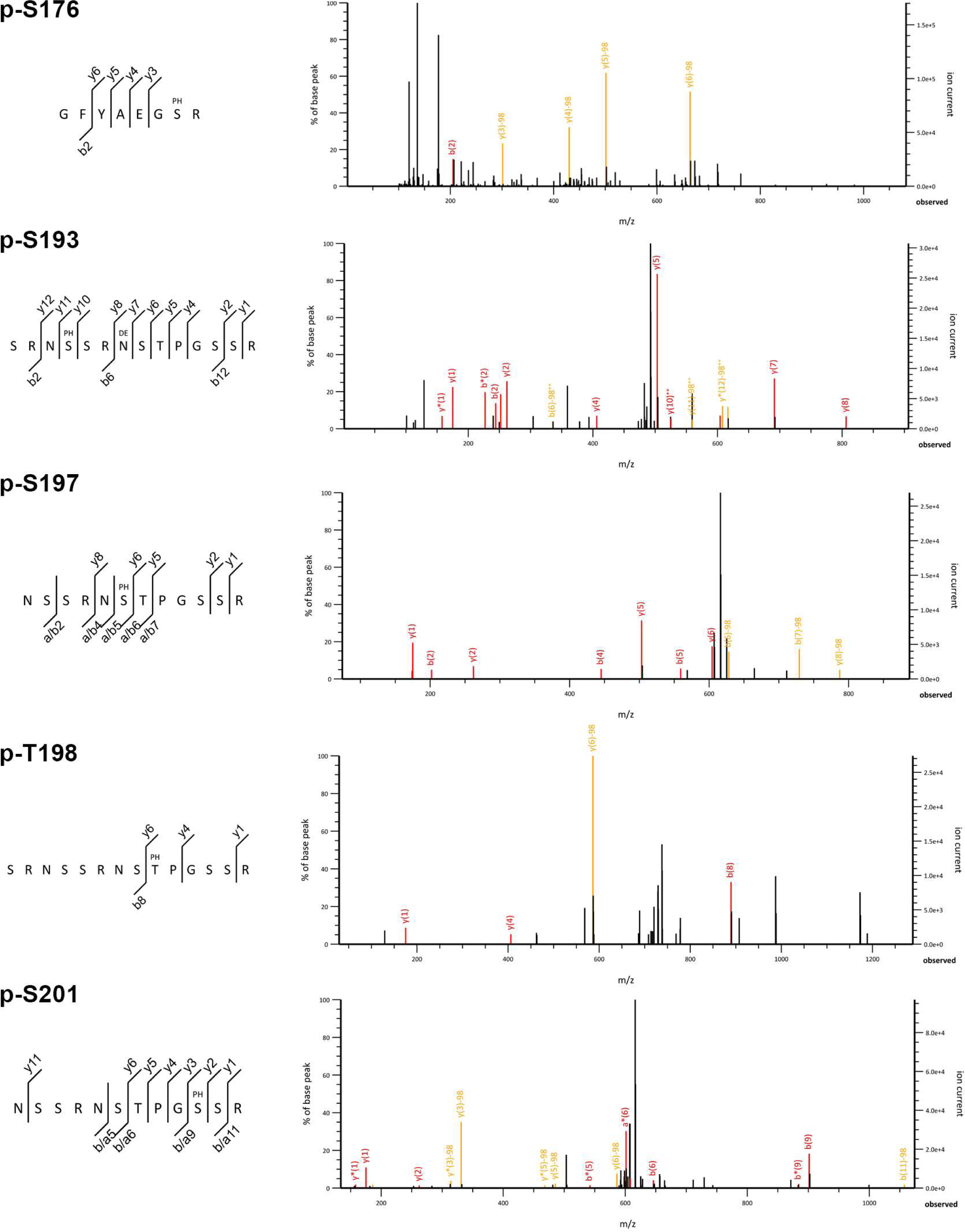
MS/MS product ion spectra of the peptide precursor ions used for mapping the phosphorylation sites within the SR domain (in addition to those shown in Fig. 2D). The phosphopeptide “GFYAEG<u>S</u>R” (p-S176) was obtained by complete tryptic digestion of the purified N* sample.

**Figure S4.**
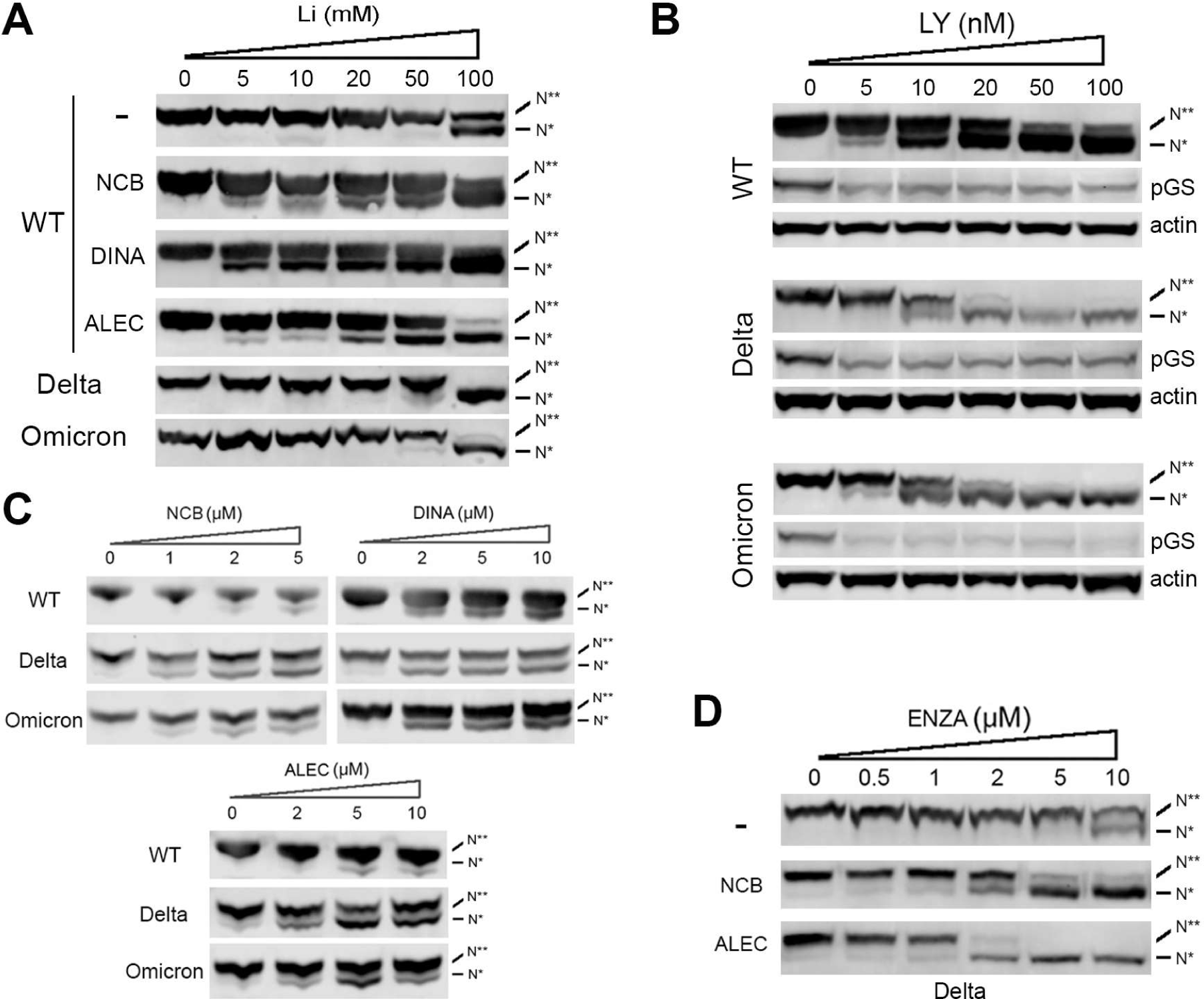
The effects of GSK-3 inhibitors [lithium, LY2090314 (LY), enzastaurin (ENZA)] and priming kinase inhibitors (NCB-0846, dinaciclib, alectinib) on N hyperphosphorylation were tested in 293T cells stably expressing the (Strep II)_2_-tagged wildtype and mutant N proteins. (**A**) The priming kinase inhibitors (1μM) increased the sensitivity to lithium. The dose-response relationship of lithium was also determined for the delta (R203M) and omicron (R203K/G204R) mutants. (**B**) A higher concentration of LY2090314 is required to inhibit the phosphorylation of N than that of an endogenous GSK-3 substrate (GS). Both delta and omicron mutants have a steeper dose-response curve (IC_50_∼10nM), whereas the WT has a shallow titration with incomplete inhibition even at 100nM LY2090314. (**C**) The effect of delta and omicron mutations on the dose-response relationship between N hyperphosphorylation and priming kinase inhibitors. (**D**) The two orally available priming kinase inhibitors (NCB-0846, alectinib), both at 2μM, increase the sensitivity of N (delta) hyperphosphorylation to enzastaurin. However, only alectinib enables enzastaurin to produce >50% inhibition at a physiologically achievable concentration of 2μM (Carducci et al., 2006).

**Figure S5.**
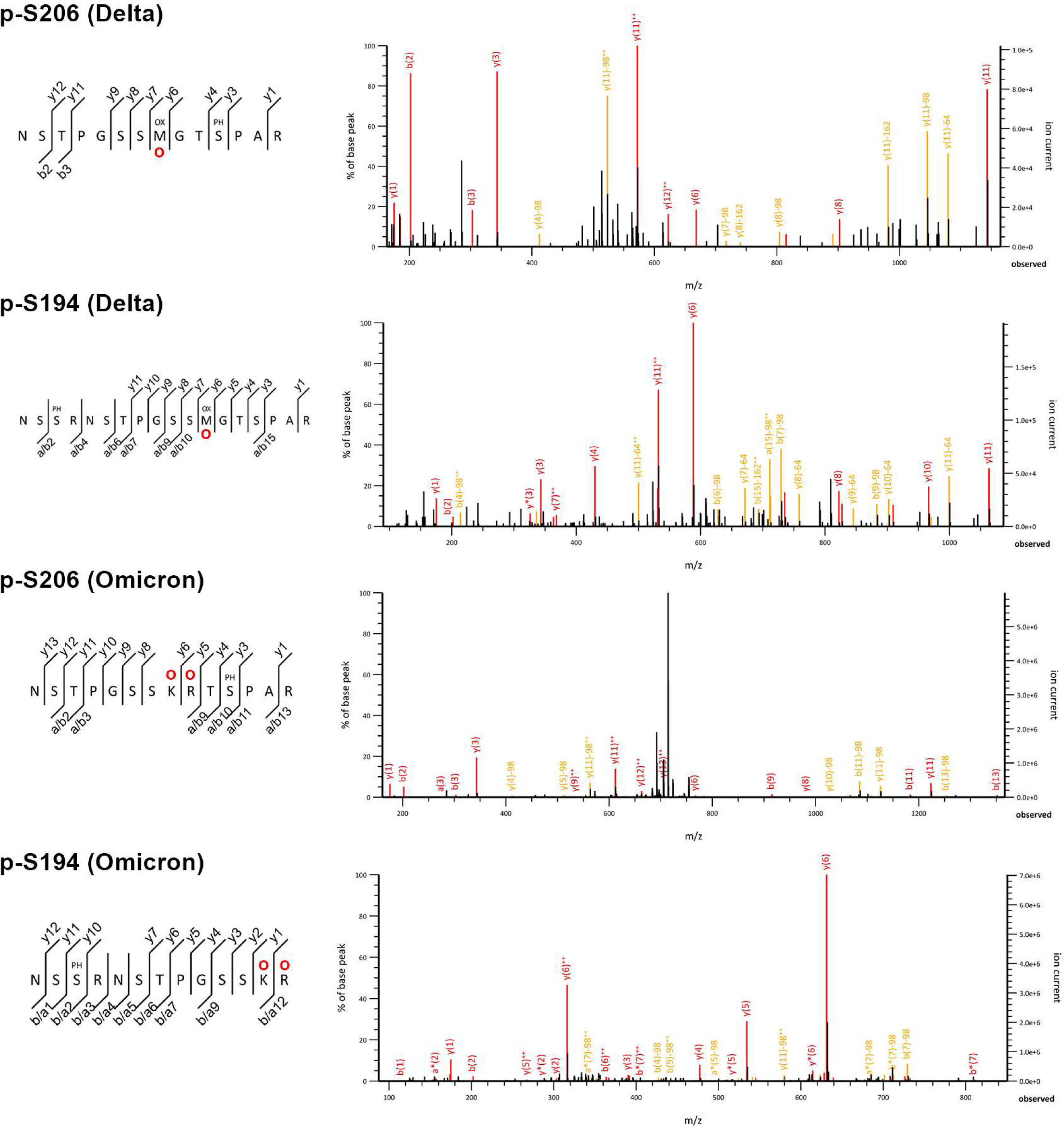
MS/MS product ion spectra of the peptide precursor ions used for mapping and quantifying the phosphorylated Ser-206 and Ser-194 in the delta and omicron mutant N proteins. The mutated amino acids are indicated by red circles.

**Figure S6.**
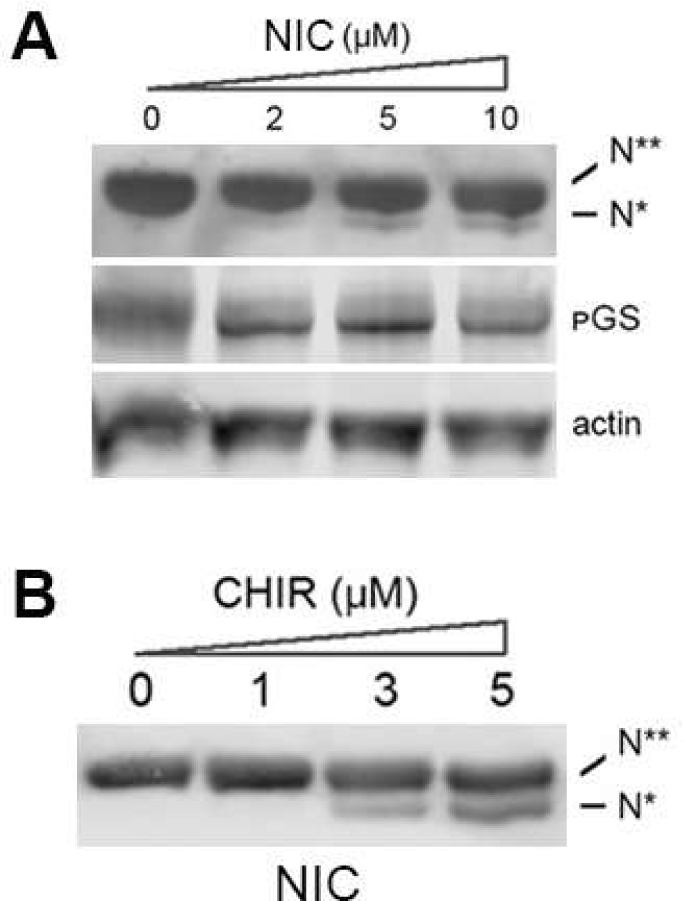
Niclosamide (NIC) weakly inhibits N hyperphosphorylation in 293T cells stably expressing the (Strep II)_2_-tagged wildtype N protein (**A**), but at a concentration of 2μM, does not significantly change the sensitivity to GSK-3 inhibitor CHIR-99021 (**B**).

---

**Table S1.** Sequence variations within the SR domain of SARS-CoV-2’s N protein. The a.a. number and type in the reference sequence (hCoV-19/Wuhan/WIV04/2019) are shown in the top two rows. The left column contains the 20 types of a.a. (“X” indicates an unknown a.a. due to uncertainty in the deposited genome sequence). The numbers inside the table indicate how many times a particular a.a. is found at that position.

|  | 171 | 172 | 173 | 174 | 175 | 176 | 177 | 178 | 179 | 180 | 181 | 182 | 183 | 184 | 185 | 186 | 187 | 188 | 189 | 190 |
| --- | --- | --- | --- | --- | --- | --- | --- | --- | --- | --- | --- | --- | --- | --- | --- | --- | --- | --- | --- | --- |
|  | F | Y | A | E | G | S | R | G | G | S | Q | A | S | S | R | S | S | S | R | S |
| A | 5 | 0 | 5816270 | 1 | 0 | 2 | 0 | 13 | 4 | 0 | 0 | 5815353 | 211 | 10 | 0 | 62 | 68 | 7 | 0 | 0 |
| C | 28 | 5 | 0 | 0 | 0 | 0 | 0 | 148 | 371 | 51 | 0 | 0 | 4 | 1 | 1162 | 1 | 0 | 0 | 245 | 23 |
| D | 1 | 5 | 0 | 3 | 0 | 0 | 0 | 541 | 449 | 0 | 0 | 90 | 0 | 0 | 0 | 0 | 0 | 0 | 0 | 1 |
| E | 0 | 0 | 2 | 5825115 | 1 | 0 | 2 | 0 | 0 | 0 | 42 | 0 | 0 | 0 | 0 | 0 | 0 | 0 | 0 | 0 |
| F | 5820150 | 4 | 0 | 0 | 0 | 0 | 0 | 2 | 0 | 0 | 0 | 10 | 111 | 46 | 1 | 690 | 1 | 0 | 0 | 0 |
| G | 1 | 0 | 0 | 1 | 5827402 | 11 | 90 | 5820736 | 5815476 | 87 | 1 | 1 | 0 | 0 | 15 | 0 | 0 | 0 | 6 | 429 |
| H | 0 | 17 | 0 | 0 | 0 | 0 | 0 | 0 | 0 | 0 | 181 | 0 | 0 | 0 | 174 | 0 | 2 | 2 | 484 | 0 |
| I | 54 | 0 | 2 | 0 | 0 | 8 | 39 | 0 | 0 | 1749 | 0 | 0 | 0 | 0 | 0 | 0 | 0 | 0 | 0 | 5465 |
| K | 0 | 0 | 0 | 2 | 0 | 0 | 113 | 0 | 0 | 0 | 48 | 0 | 0 | 0 | 0 | 0 | 3 | 0 | 0 | 0 |
| L | 97 | 0 | 0 | 0 | 0 | 0 | 0 | 0 | 0 | 0 | 690 | 0 | 2 | 5 | 256 | 0 | 15410 | 1249 | 319 | 0 |
| M | 0 | 0 | 0 | 0 | 0 | 0 | 0 | 0 | 0 | 0 | 0 | 0 | 0 | 0 | 0 | 0 | 0 | 0 | 0 | 0 |
| N | 0 | 3 | 0 | 0 | 0 | 4 | 0 | 0 | 0 | 170 | 0 | 0 | 0 | 0 | 0 | 2 | 1 | 0 | 0 | 596 |
| P | 0 | 0 | 3 | 0 | 0 | 0 | 0 | 0 | 0 | 0 | 6 | 86 | 642 | 62 | 190 | 168 | 180 | 159 | 3 | 0 |
| Q | 0 | 0 | 0 | 6 | 0 | 0 | 0 | 0 | 0 | 0 | 5821673 | 1 | 0 | 0 | 0 | 0 | 0 | 0 | 0 | 0 |
| R | 0 | 0 | 0 | 0 | 15 | 14 | 5825317 | 40 | 13 | 43 | 126 | 0 | 0 | 0 | 5821819 | 1 | 0 | 0 | 5820293 | 819 |
| S | 3 | 1 | 1067 | 0 | 0 | 5828314 | 17 | 465 | 3826 | 5818731 | 1 | 7364 | 5810571 | 5824282 | 30 | 5821278 | 5808513 | 5821520 | 69 | 5813483 |
| T | 0 | 0 | 103 | 1 | 1 | 3 | 0 | 0 | 0 | 419 | 1 | 273 | 4440 | 4 | 0 | 16 | 30 | 3 | 0 | 2 |
| V | 10 | 0 | 4269 | 1 | 4 | 0 | 0 | 925 | 1641 | 2 | 0 | 325 | 0 | 0 | 2 | 0 | 0 | 0 | 2 | 3 |
| W | 0 | 0 | 0 | 0 | 3 | 0 | 0 | 0 | 0 | 0 | 0 | 0 | 2 | 0 | 0 | 0 | 0 | 0 | 0 | 0 |
| Y | 42 | 5819825 | 1 | 0 | 0 | 0 | 0 | 0 | 0 | 0 | 0 | 5 | 7055 | 16 | 0 | 1477 | 0 | 0 | 1 | 0 |
| X | 51261 | 51792 | 49935 | 46522 | 44226 | 43296 | 46074 | 48782 | 49872 | 50400 | 48883 | 48144 | 48614 | 47226 | 48003 | 47957 | 47444 | 48712 | 50230 | 50831 |

|  | 191 | 192 | 193 | 194 | 195 | 196 | 197 | 198 | 199 | 200 | 201 | 202 | 203 | 204 | 205 | 206 | 207 | 208 | 209 | 210 |
| --- | --- | --- | --- | --- | --- | --- | --- | --- | --- | --- | --- | --- | --- | --- | --- | --- | --- | --- | --- | --- |
|  | R | N | S | S | R | N | S | T | P | G | S | S | R | G | T | S | P | A | R | M |
| A | 2 | 0 | 0 | 835 | 0 | 0 | 18 | 61 | 2 | 9 | 2 | 1 | 0 | 737 | 118 | 102 | 12 | 5787824 | 1 | 1 |
| C | 273 | 0 | 99 | 0 | 0 | 0 | 0 | 0 | 0 | 3051 | 192 | 1668 | 0 | 0 | 0 | 44 | 0 | 1 | 0 | 0 |
| D | 0 | 170 | 0 | 0 | 0 | 76 | 0 | 0 | 0 | 686 | 0 | 0 | 0 | 0 | 0 | 1 | 0 | 130 | 0 | 0 |
| E | 0 | 0 | 0 | 0 | 2 | 0 | 0 | 0 | 0 | 0 | 0 | 0 | 3201 | 205 | 0 | 0 | 0 | 0 | 2 | 1 |
| F | 0 | 0 | 0 | 0 | 0 | 0 | 1 | 0 | 0 | 0 | 0 | 82 | 0 | 0 | 0 | 5693 | 4 | 10 | 0 | 0 |
| G | 4 | 0 | 218 | 0 | 100 | 2 | 0 | 0 | 0 | 5822394 | 829 | 483 | 46 | 4295430 | 1 | 1 | 0 | 657 | 93 | 1 |
| H | 1112 | 7 | 0 | 0 | 0 | 151 | 0 | 0 | 1 | 0 | 0 | 0 | 0 | 0 | 0 | 0 | 135 | 0 | 0 | 0 |
| I | 0 | 3 | 10196 | 0 | 6937 | 524 | 0 | 1695 | 2 | 0 | 2933 | 13478 | 2073 | 0 | 157683 | 0 | 0 | 0 | 10689 | 11642 |
| K | 0 | 185 | 0 | 1 | 1513 | 54 | 0 | 0 | 0 | 0 | 0 | 10 | 1510292 | 1 | 1 | 0 | 0 | 0 | 1760 | 114 |
| L | 696 | 0 | 0 | 98816 | 129 | 1 | 7903 | 3 | 165571 | 0 | 0 | 0 | 40 | 2902 | 3 | 142 | 5378 | 2 | 0 | 257 |
| M | 0 | 0 | 0 | 0 | 0 | 0 | 0 | 0 | 0 | 0 | 0 | 3 | 3394776 | 1 | 2 | 0 | 0 | 0 | 1 | 5811059 |
| N | 0 | 5822858 | 592 | 0 | 10 | 5825600 | 1 | 189 | 0 | 1 | 504 | 5490 | 15 | 0 | 483 | 0 | 2 | 0 | 1 | 1 |
| P | 7 | 0 | 0 | 299 | 0 | 0 | 555 | 12 | 5648757 | 1 | 0 | 14 | 0 | 82716 | 45 | 1018 | 5810146 | 243 | 0 | 0 |
| Q | 0 | 0 | 0 | 2 | 0 | 0 | 2 | 0 | 257 | 0 | 0 | 0 | 0 | 318 | 0 | 0 | 26 | 0 | 0 | 0 |
| R | 5818934 | 1 | 185 | 0 | 5817236 | 1 | 0 | 0 | 11 | 7 | 138 | 13370 | 895037 | 1427728 | 0 | 0 | 75 | 0 | 5809749 | 80 |
| S | 74 | 115 | 5811142 | 5724498 | 230 | 701 | 5817037 | 9 | 7080 | 540 | 5821148 | 5784797 | 269 | 14 | 214 | 5821317 | 11119 | 34996 | 314 | 0 |
| T | 0 | 14 | 1913 | 15 | 1003 | 55 | 1082 | 5826867 | 277 | 0 | 0 | 106 | 762 | 1 | 5667580 | 39 | 564 | 213 | 813 | 663 |
| V | 0 | 0 | 3 | 2 | 0 | 0 | 1 | 0 | 2 | 187 | 0 | 4 | 1140 | 711 | 5 | 0 | 1 | 2847 | 0 | 283 |
| W | 0 | 0 | 0 | 0 | 0 | 0 | 0 | 0 | 0 | 0 | 0 | 0 | 4 | 0 | 0 | 0 | 0 | 0 | 0 | 2 |
| Y | 0 | 207 | 0 | 0 | 0 | 66 | 0 | 0 | 0 | 0 | 0 | 17 | 0 | 0 | 0 | 126 | 0 | 0 | 0 | 0 |
| X | 50550 | 48092 | 47304 | 47184 | 44492 | 44421 | 45052 | 42816 | 49692 | 44776 | 45906 | 52129 | 63997 | 60888 | 45517 | 43169 | 44190 | 44729 | 48229 | 47548 |

**Table S2.**
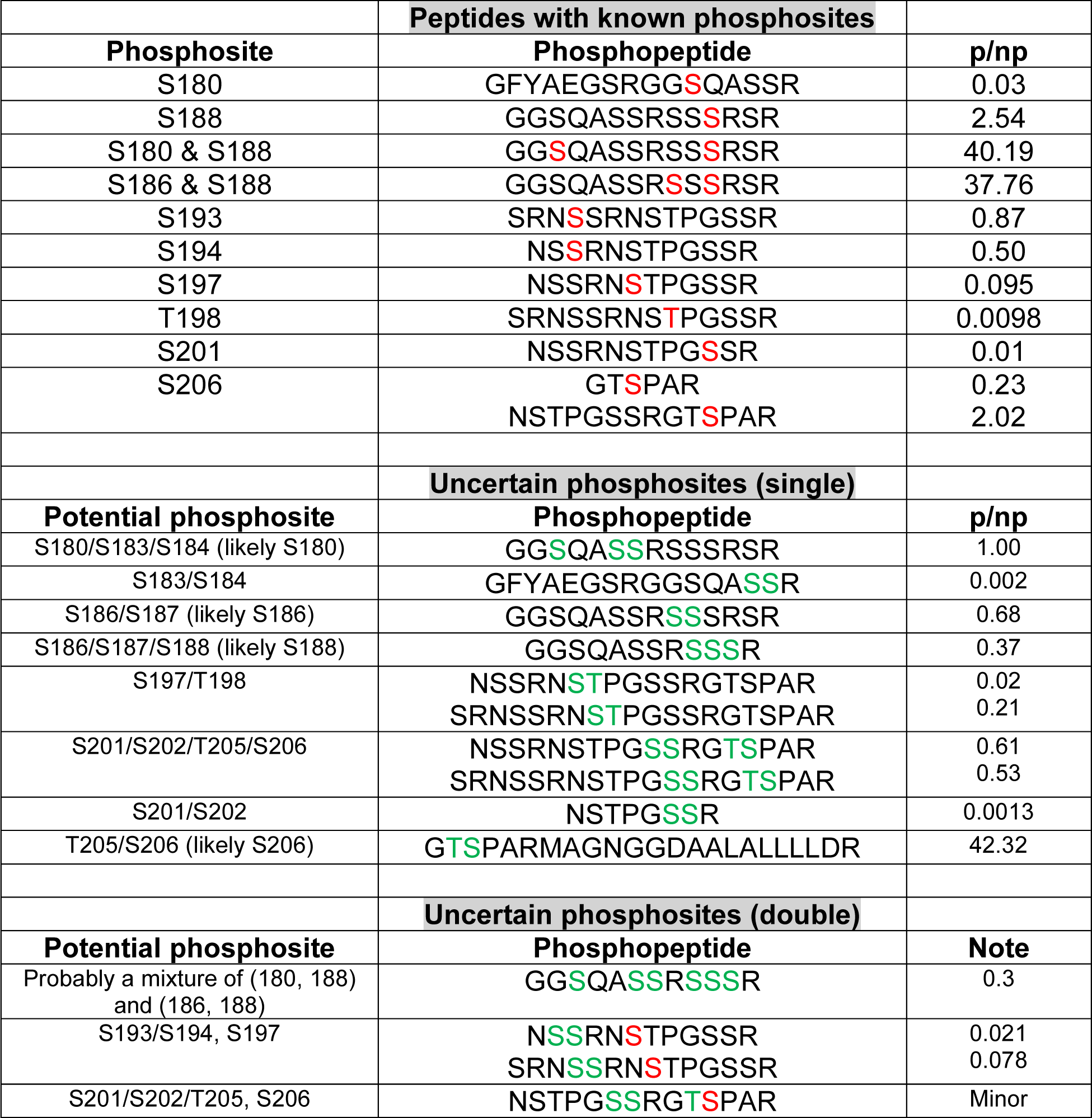
GSK3-independent phosphorylation sites within the SR domain. A total of 14 singly or doubly phosphorylated peptides, covering the entire SR domain, were observed. From these, 9 phosphosites (highlighted in red) could be unequivocally mapped based on the MS/MS spectrum (in the lower portion of the table, likely phosphorylation sites are highlighted in green). The relative abundance (phosphorylation stoichiometry) of the known site(s) could be estimated by comparing the areas under the LC peaks for the phosphorylated peptide (p) and its unphosphorylated counterpart (np), assuming the two peptides have similar ionization efficiency. In a separate experiment where N was thoroughly digested by trypsin, we also observed a phosphopeptide containing p-S176.

